# IL-10 suppresses T cell expansion while promoting tissue-resident memory cell formation during SARS-CoV-2 infection in rhesus macaques

**DOI:** 10.1101/2022.09.13.507852

**Authors:** Christine E. Nelson, Taylor W. Foreman, Keith D. Kauffman, Shunsuke Sakai, Sydnee T. Gould, Joel D. Fleegle, Felipe Gomez, NIAID/DIR Tuberculosis Imaging Program, Cyril Le Nouën, Xueqiao Liu, Tracey L. Burdette, Nicole L. Garza, Bernard A. P. Lafont, Kelsie Brooks, Cecilia S. Lindestam Arlehamn, Daniela Weiskopf, Alessandro Sette, Heather D. Hickman, Ursula J. Buchholz, Reed F. Johnson, Jason M. Brenchley, Laura E. Via, Daniel L. Barber

## Abstract

The pro- and anti-inflammatory pathways that determine the balance of inflammation and viral control during SARS-CoV-2 infection are not well understood. Here we examine the roles of IFNγ and IL-10 in regulating inflammation, immune cell responses and viral replication during SARS-CoV-2 infection of rhesus macaques. IFNγ blockade tended to decrease lung inflammation based on ^18^FDG-PET/CT imaging but had no major impact on innate lymphocytes, neutralizing antibodies, or antigen-specific T cells. In contrast, IL-10 blockade transiently increased lung inflammation and enhanced accumulation of virus-specific T cells in the lower airways. However, IL-10 blockade also inhibited the differentiation of virus-specific T cells into airway CD69^+^CD103^+^ T_RM_ cells. While virus-specific T cells were undetectable in the nasal mucosa of all groups, IL-10 blockade similarly reduced the frequency of total T_RM_ cells in the nasal mucosa. Neither cytokine blockade substantially affected viral load and infection ultimately resolved. Thus, in the macaque model of mild COVID-19, the pro- and anti-inflammatory effects of IFNγ and IL-10 have no major role in control of viral replication. However, IL-10 has a key role in suppressing the accumulation of SARS-CoV-2-specific T cells in the lower airways, while also promoting T_RM_ at respiratory mucosal surfaces.

## INTRODUCTION

SARS-CoV-2 infection has a spectrum of clinical disease outcomes, ranging from asymptomatic to fatal. The severity of COVID-19, the disease caused by SARS-CoV-2 infection, is determined by the balance of viral replication and immune-mediated pathology. The factors that prevent or promote pulmonary inflammation during SARS-CoV-2 infection, however, are not well understood. The majority of experimentally SARS-CoV-2-infected rhesus macaques develop only mild symptoms and rapidly clear virus (*1, 2*). Accordingly, this species may be a useful model for examining the mechanisms of effective viral control and limited inflammatory disease that occurs in most humans after infection with SARS-CoV-2. Here we use rhesus macaques to examine the roles of IFNγ and IL-10, prototypic pro- and anti-inflammatory cytokines respectively, in host resistance to SARS-CoV-2 infection and the development of COVID-19 disease.

IFNγ and molecules induced by IFNγR signaling (e.g., CXCL10) have been associated with severe COVID-19 and the development of acute respiratory distress syndrome (*3-12*). Elevated levels of IFNγ also strongly correlate with the development of multi-system inflammatory syndrome in children after SARS-CoV-2 infection (*8, 13, 14*). In ACE2 transgenic mice, neutralizing IFNγ along with TNF reduced mortality of severe SARS-CoV-2 infection (*7, 15*). Therefore, IFNγ driven inflammation may contribute to lung pathology during severe COVID-19. Furthermore, the addition of IFNγ to an organoid cell culture system, IFNγ increased ACE2 expression and enhanced SARS-CoV-2 replication (*16*). Despite these potential detrimental effects of IFNγ, it remains possible that IFNγ may also contribute to host-protection in some settings. Indeed, the role of IFNγ in vivo during SARS-CoV-2 infection is not well explored.

IL-10 has also been associated with severe COVID-19 (*5, 17, 18*). IL-10 is upregulated early in disease progression, and along with IL-6, is a predictive biomarker for poor COVID-19 outcomes (*5, 17*). However, in children, one study reported that higher plasma IL-10 levels were correlated with decreased viral measurements from nasal aspirates (*19*). Therefore, the precise role of IL-10 during SARS-CoV-2 infection and COVID-19 pathogenesis *in vivo* is not well defined.

Studies have shown that SARS-CoV-2 replicates in the upper and lower airways of rhesus macaques and an early wave of localized lung inflammation correlates with viral clearance (*1, 2*). In the present study, animals were treated with IFNγ and IL-10 blocking reagents beginning the day before infection with SARS-CoV-2. Using longitudinal ^18^FDG-PET/CT imaging we quantified SARS-CoV-2 induced inflammation in the chest and head. We also measured viral replication, innate lymphocyte activation, and Ag-specific T and B cell responses after cytokine blockade. We find that IL-10 and IFNγ counter-regulate lung inflammation without an appreciable effect on viral replication. Moreover, we identify a key role for IL-10 in negatively regulating the clonal expansion of virus-specific T cells while promoting T_RM_ cells in respiratory mucosal surfaces.

## RESULTS

### ^18^FDG-PET/CT analysis of lung inflammation

To investigate the role of pro- and anti-inflammatory cytokines in viral replication and pathogenesis during SARS-CoV-2 infection, we treated 15 male rhesus macaques with either a primatized monoclonal antibody targeting IL-10 (anti-IL-10), a rhesus macaque IFNγR1-immunoglobulin fusion protein (rmIFNγR1-Ig), or isotype control (anti-DSP IgG1) by the intravenous (i.v.) route, with five animals per group (Fig. 1A). One day after treatment, animals were infected with a total of 2×10^6^ TCID_50_ of SARS-CoV-2/USA-WA-1 split between the intranasal (i.n.) and intratracheal (i.t.) routes. An additional dose of blocking reagent was given at day 3 post-infection. Using IL-10 and IFNγ reporter cell lines, we verified that anti-IL-10 and rmIFNγR1-Ig efficiently blocked IL-10 and IFNγ signaling respectively, *in vitro* (Fig. S1). We monitored inflammation by ^18^Flurorine deoxyglucose (^18^FDG)-positron emission tomography/computed tomography (PET/CT) imaging of the head, chest, and abdomen. In control animals, we observed evidence of lung inflammation with increased density and ^18^FDG-avidity that peaked at day 2 and resolved by days 6-10 post-infection, which is consistent with previous findings (Fig. 1B-D, S2) (*1, 20, 21*). Animals receiving IL-10 blocking antibody had an increased number of lesions that were more metabolically active on day 6 post-infection, as compared to isotype controls or rmIFNγR1-Ig treated animals (Fig. 1C, D). Each lesion was given an intensity score based on the sum of the normalized maximum values for lesion size, FDG uptake, and density in Hounsfield’s units (HU) (Fig. 1E). Lesions from anti-IL-10 treated animals had increased lesion intensity scores compared to controls. At day 6 post-infection when most of the inflammation had resolved in control animals, lesions in the anti-IL-10 treated animals still had significant ^18^FDG uptake and some lesions had increased in intensity from that observed at day 2 post-infection (Fig. 1D). Conversely, animals that received rmIFNγR1-Ig tended to have decreased number, size, and density of lung lesions as compared to isotype controls or anti-IL-10 treated (Fig. 1C-E, S2). In contrast to the isotype control and anti-IL-10 treated, the few lesions that were present in the rmIFNγR1-Ig treated animals were also significantly less dense and metabolically active at day 6 compared to day 2 (Fig. 1D).

**Figure 1.**
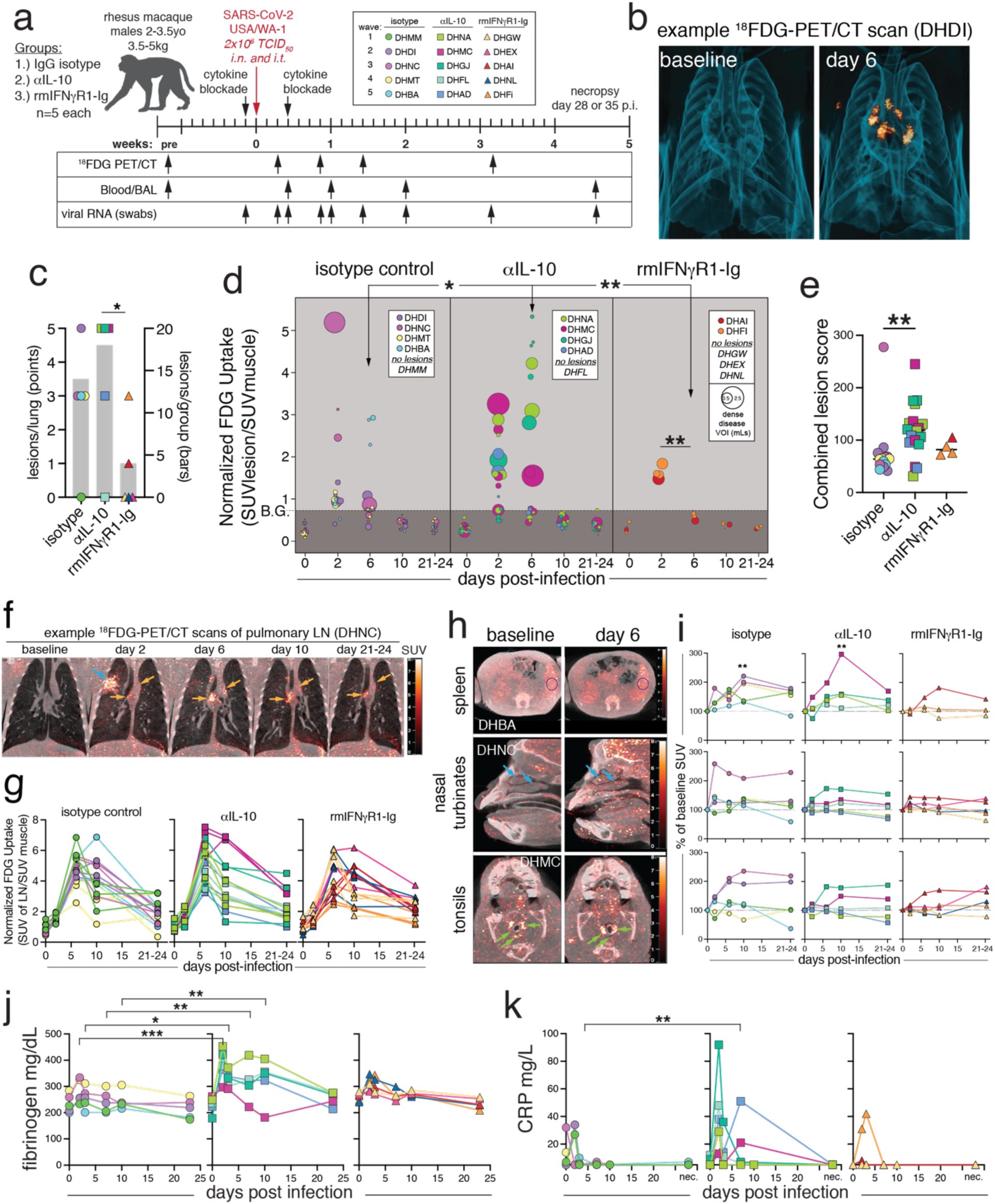
SARS-CoV-2 induced lung inflammation is increased with IL-10 blockade and decreased with IFNγ blockade. (A) Experimental design: Fifteen male rhesus macaques in three treatment groups: IgG isotype control, anti-IL-10, or anti-IFNγ (rmIFNγR1-Ig), with n=5 per group. Animals were treated with 10mg/kg of monoclonal antibody i.v. one day prior to infection and three days after infections with SARS-CoV-2/USA/WA-1 at a dose of 2×10^6^ TCID_50_, administered intranasal (i.n.) and intratracheal (i.t.). (B) 3D rendering of representative lung ^18^FDG-PET/CT images from baseline and 6 post infection from isotype control animal DHDI. (C) Number of lesions per animal (left axis, points) and average number of lesions per group (right axis, grey bars). Significance calculated with individual t-test with Welch’s correction for lesions per animal. (D) Quantification of FDG uptake in standard uptake value (SUV) normalized to muscle, and volume of individual lesions (size of dot), based on volume of interest (VOI) > −550 Hounsfield units (HU) defined at days 2 or 6 post-infection. Significance calculated with individual t-test with Welch’s correction for FDG uptake at day 6 between groups and Tukey’s multiple comparison test of day 2 vs. day 6 within each group. (E) Lesion score for individual lesions calculated as the sum of normalized max FDG uptake, normalized max Hounsfield’s units, and normalized max volume. Significance calculated with Dunn’s multiple comparison test. (F) Example PET/CT images showing pulmonary lymph node FDG signal from baseline, day 2, 6, 10, and 21-24 post infection from isotype control animal DHNC. Orange arrows indicate lymph nodes and blue arrows indicate lung lesions. (G) Quantification of metabolic activity of lymph nodes as measured FDG uptake in SUV, normalized to muscle. All timepoints post-infection were statistically significant over baseline by 2-way ANOVA and Tukey’s multiple comparison test. (H) Example PET/CT images with evidence of FDG signal from spleen, nasal turbinates, and tonsils from baseline and day 6 post infection. Animal IDs are embedded in image. DHBA and DHNC (isotype). DHMC (anti-IL10). (I) Quantification of change in FDG uptake (SUV) calculated as change from baseline for each animal with detectable signal from spleen, nasal turbinates, and tonsils. Significance calculated by 2-way ANOVA and Tukey’s multiple comparison test. (J) Plasma fibrinogen levels in mg/dL. Significance calculated with 2-way ANOVA and a Dunnett’s multiple comparison test. (K) Plasma C-reactive protein (CRP) in mg/L. Limit of detection >5mg/L. Significance calculated with a 2-way ANOVA and a Dunnett’s multiple comparison test.

### ^18^FDG-PET/CT analysis of extra-pulmonary inflammation

^18^FDG uptake in the pulmonary lymph nodes (pLN) began to increase by day 2 post-infection and peaked at day 6-10 post-infection, with some lymph nodes retaining ^18^FDG avidity on day 22-23 post-infection (Fig. 1F, G, S2). Unlike the lungs, there were no differences between treatment groups in the PET/CT signal from pLNs after infection (Fig. 1G). Comparing the percent of maximum FDG uptake in the lymph nodes versus the lungs revealed a significant difference in the kinetics of inflammation between the two compartments. The lymph nodes had delayed but prolonged PET/CT signal as compared to the lung, perhaps reflecting the kinetics of ongoing immune responses in the lung draining lymph nodes (Fig. S2C).

**Figure 2.**
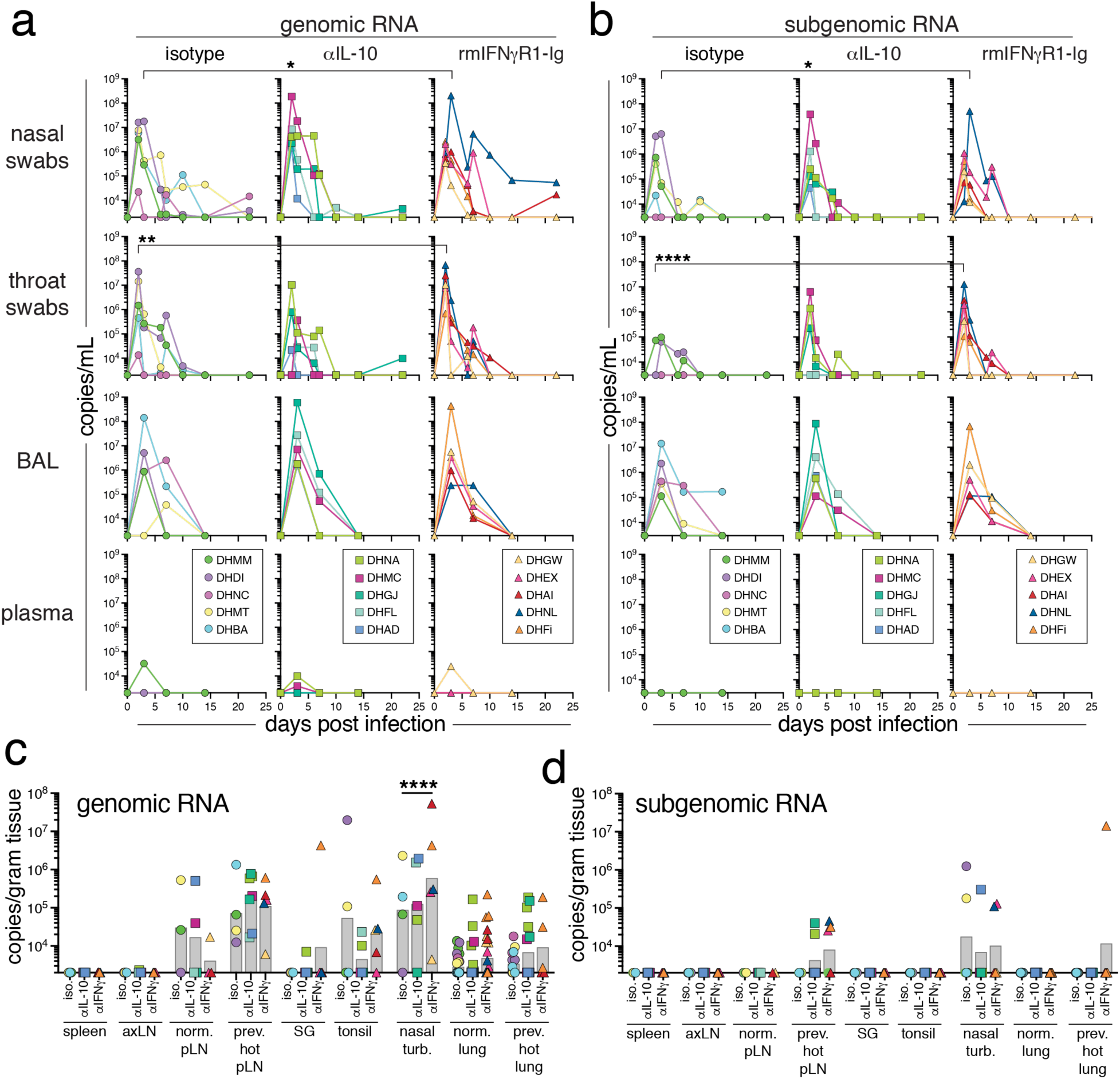
IFNγ and IL-10 are not required for suppression of SARS-CoV-2 replication. (A) Genomic and (B) Subgenomic RNA quantification of the N gene of SARS-CoV-2 by RT-qPCR in copies/mL from bronchoalveolar lavage (BAL), nasal swabs, throat swabs, and plasma. (C) Genomic and (D) Subgenomic RNA quantification of the N gene by RT-qPCR in copies/gram of tissue by RT-qPCR at necropsy (day 28-35 post-infection) from spleen, axillary lymph node (axLN), normal pulmonary lymph nodes (norm. pLN), previously PET/CT hot pulmonary lymph nodes (prev. hot pLN), salivary gland (SG), tonsil, nasal turbinates (nasal turb.), normal lung sections (norm. lung), and previously PET/CT hot lung sections. For *A* and *B* the cutoff for RNA detection is 2,000 copies/mL for genomic RNA and 3,000 copies/mL for subgenomic RNA. For *C* and *D* the cutoff is 2,000 copies/gram of tissue for both genomic and subgenomic RNA. Graphs show individual animals from samples taken at baseline, days 2, 3, 6, 7, 10, 14, and 22 post-infection, as well as necropsy. Significance calculated with a 2-way ANOVA and a Dunnett’s multiple comparison test

We further investigated SARS-CoV-2 induced extra-pulmonary inflammation using ^18^FDG-PET/CT analysis of the spleen, nasal turbinates, and tonsils. There was increased ^18^FDG uptake in the spleens of isotype control and anti-IL-10 treated animals, that peaked at ∼day 10 post-infection and reached statistical significance over baseline (Fig. 1H, I). However, only 1 of 5 rmIFNγR1-Ig treated animals had increased ^18^FDG uptake in the spleen above baseline at day 10, which did not reach statistical significance. The nasal turbinates also had evidence of modest ^18^FDG avidity that peaked at ∼day 2-6 post-infection. However, these changes did not reach statistical significance over baseline, and there were no differences between treatment groups. Lastly, low levels of ^18^FDG uptake were observed in the tonsils of in some animals at ∼day 10 post-infection. However, no significance differences were observed between groups. Collectively, these data suggest that the lung is the primary site of inflammation following SARS-CoV-2 infection of rhesus macaques. Moreover, IL-10 and to a lesser extent IFNγ modulated pulmonary but not extrapulmonary inflammation in this model.

### Clinical measures of COVID-19 disease severity

Clinical measurements were assessed after SARS-CoV-2 infection including body weight, temperature, oxygen saturation (SpO_2_), heart rate, respiratory rate, fasting blood glucose, and plasma levels of C-reactive protein (CRP), D-dimer, and fibrinogen (Fig. 1J, K, S3). Plasma fibrinogen levels were higher in the anti-IL-10 treated animals compared to controls at days 2-10 post-infection, suggestive of increased coagulation (Fig. 1J). CRP levels were also elevated in the anti-IL-10 treated animals at day 2 post-infection, as compared to controls (Fig. 1K). There was a decrease in D-dimer levels in the rmIFNγR1-Ig treated animals at day 3 post-infection that reached statistical significance over controls, but this was difficult to interpret given the variability at baseline (Fig. S3F). There was a noted increase in fasting blood glucose levels at day 2 after SARS-CoV-2 infection that reached statistical significance over baseline in the anti-IL-10 and anti-IFNγ treated groups; however, these differences were not significantly increased over controls (Fig. S3E). There was no evidence of fever, tachycardia, bradycardia, hyperventilation, or weight loss greater than 5 percent of baseline in any of the groups, consistent with overall mild disease in this model (Fig. S3). The observed differences were consistent with increased lung inflammation after IL-10 blockade and decreased inflammation with IFNγ blockade after SARS-CoV-2 infection.

**Figure 3.**
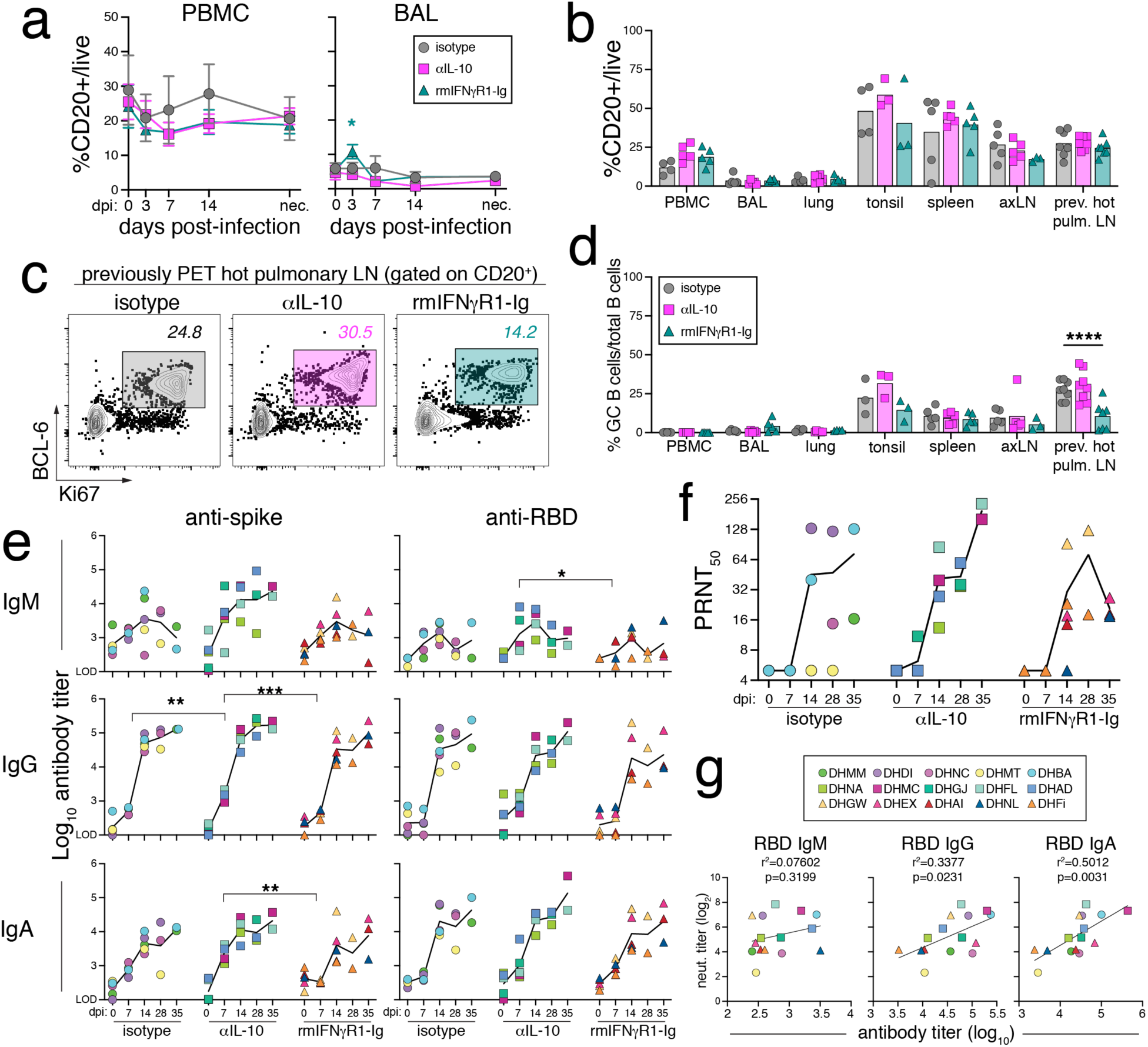
B cell and antibody are minimally impacted by IL-10 and IFNγ blockade. (A) Frequency of CD20^+^ B cells in PBMCs and BAL as a percentage of total live lymphocytes at baseline and days 3, 7, 14, and 28/35 post-infection necropsy (nec.). Lines are mean and error bars are SEM. (B) Frequency of total B cells from PBMCs, BAL, lung, tonsil, spleen, axillary lymph node (axLN), and previously hot pulmonary lymph nodes (prev. hot pulm. LN) as a percentage of total live lymphocytes at 28/35 post-infection necropsy. (C) Representative flow cytometry plots of gating strategy for germinal center B cells from animals DHDI, DHGJ, and DHEX isolated from previously PET hot pulmonary lymph nodes. Gates in plots show frequency of germinal center B cells as a frequency of CD20^+^ total B cells. (D) Quantification of germinal center B cells as a frequency of total B cells from the indicated tissues as necropsy. (E) Plasma anti-spike (left graphs) and anti-RBD (right graphs) antibody titers of the indicated isotypes: IgM, IgA, and IgG, in log_10_ endpoint titer at baseline and days 3, 7, 14, and 28 or 35 post-infection necropsy. Each animal is represented as a point and the mean as a line for each treatment group. Significance calculated with 2-way ANOVA and Tukey’s multiple comparison test. Significance not reported for comparisons with <3 data points. (F) Live-virus plasma neutralization titer at baseline and days 3, 7, 14, and 28 or 35 post-infection necropsy. Titer calculated as a 50 percent plaque reduction neutralization titer (PRNT_50_) of negative control sera. Each animal is represented as a point and the mean as a line for each treatment group. (G) Correlation analysis of neutralization titer and anti-RBD IgM, IgG, or IgA titer at 28 or 35 post-infection necropsy using simple linear regression with r-squared of goodness of fit and p-value of non-zero slope reported.

### SARS-CoV-2 replication kinetics and tissue distribution

To determine whether IL-10 or IFNγ blockade modulated SARS-CoV-2 replication, we measured genomic and subgenomic RNA of the nucleocapsid (N) gene in bronchoalveolar lavage fluid (BAL), plasma, nasal swabs, and throat swabs. Genomic (gN) and subgenomic (sgN) viral RNA levels had similar dynamics after infection, although copies of sgN were generally ∼2 logs lower compared to gN. In the BAL, viral RNA levels were highest at day 3 post-infection and fell below the limit of detection (L.O.D.) by days 7-14, with no differences detected between the groups (Fig. 2A, B). In nasal and throat swabs, gN and sgN levels peaked at ∼2-3 days post-infection and were undetectable in most animals by ∼14 days post-infection (Fig. 2A, B). There was a statistically significant increase in viral RNA in the nasal and throat swabs on day 2-3 post infection in the rmIFNγR1-Ig treated animals compared to controls; however, this difference was largely driven by the high RNA loads in animal *DHNL.* We did not observe any statistical difference in viral RNA loads in the swabs after IL-10 blockade. There was very little viral RNA detected in plasma samples, i.e., only 4 of 15 animals had detectable gN in plasma at day 3 post-infection, and there was no difference between the groups.

At necropsy (day 28 or 35 post-infection), no viral RNA was quantified above the L.O.D. in the spleen or axillary lymph nodes (axLN) (Fig. 2C, D). However, gN was still present in the pLNs, with a greater number of the previously PET-hot pLN having detectable gN compared to uninvolved pLN (normal pLN). Subgenomic RNA was also detected in the previously PET-hot pLN but was below the L.O.D. in the normal pLN from all animals. There were no differences in lymph node viral RNA levels between groups.

At necropsy, non-lymphoid tissues including the salivary gland (SG), tonsils, nasal turbinates, and lung, were also assessed for viral RNA loads (Fig. 2C, D). Genomic RNA was detected in only two SG samples, and no sgN was detected in any SG samples. Tonsils contained low levels of gN in 8 of 14 samples tested, but no sgN was detected and there was no difference between groups. Nasal turbinates had higher levels of viral RNA compared to SG and tonsils. Like the nasal and throat swabs, there was a statistically significant increase in gN detected of the nasal turbinates in the rmIFNγR1-Ig treated animals compared to controls; although, no difference was found in sgN between groups (Fig. 2C). Lung sections from previously PET-hot regions were isolated separately from uninvolved lung sections (normal lung), as described previously (*1*). In the normal lung sections, gN was detected in 9 of 29 isotype control, 5 of 30 in anti-IL-10 treated, and 12 of 30 in rmIFNγR1-Ig treated sections (Fig. 2C). In previously PET-hot lung sections, gN was detected in 7 of 15 control, 6 of 16 anti-IL-10 treated, and 3 of 5 rmIFNγR1-Ig treated samples, and there was no statistical difference between the treatment groups. Subgenomic RNA was not detected in any normal lung sections, and in only one of the previously PET-hot lung sections isolated from an rmIFNγR1-Ig treated animal (Fig. 3D). Altogether, there was little impact of either cytokine blockade on SARS-CoV-2 viral RNA loads, although IFNγ blockade may have transiently increased viral replication in the upper airway early during infection.

### NK and MAIT cell responses

NK cells are important in early control of viral infections and have been shown to produce IFNγ in response to SARS-CoV-2 (*22*). Studies of COVID-19 patients suggest that NK cells might become dysfunctional during SARS-CoV-2 infection (*22-25*). Therefore, we investigated NK cells response in the peripheral blood mononuclear cell (PBMC) compartment and BAL. In NHP, NK cells can be defined as CD3^-^/CD8α^+^/CD8α^-^ /NKG2A^+^ and can be divided into 4 distinct subpopulations based on the expression of CD16 and CD56 (*26-29*). The population of CD16^+^/CD56^-^ NK cells have been shown to be cytotoxic, with the expression of perforin and granzyme B (*26*). Whereas CD16^-^/CD56^+^ NK cells are thought to be less cytotoxic and more likely to produce IFNγ (*28*). In PBMCs of rhesus macaques prior to infection, we observed a skewing of the NK population to a predominately CD16^+^/CD56^-^ cytotoxic phenotype, which is consistent with previous reports (Fig. S4A, B) (*27*). In the BAL, we observed an increase in the proportion of CD16^-^/CD56^+^ cytokine producing NK cells. We also observed a greater abundance of CD16^+^/CD56^+^ intermediate and CD16^-^/CD56^-^ populations in the BAL, as compared to PBMCs. After SARS-CoV-2 infection, there was an increase in total NK cells beginning at day 3 post-infection that peaked at day 7 in isotype control samples in both the PBMCs and BAL (Fig. S4B). All NK cell subsets in PMBCs had upregulated Ki67, a marker of cell-cycle/proliferation, by day 7 post-infection (Fig. S4C, D). Ki67 upregulation was less robust in the BAL than in PBMCs, especially for the CD16^+^/CD56^-^ NK cell population, which only reached significant increase over baseline at day 7 post-infection in the anti-IL-10 group (Fig. S4D).

We detected differences in the expression of granzyme B by NK cell subsets, with CD16^+^/CD56^-^ in both the PBMC and BAL having the highest expression of granzyme B at baseline (Fig. S4E, F). The population of CD16^+^/CD56^+^ in PBMCs also had high expression of granzyme B, but this population was rare in PBMCs. All NK subsets in both PBMC and BAL upregulated granzyme B at day 3 in response to SARS-CoV-2 infection. At day 7 post infection, the granzyme B response remained comparable to day 3 or was beginning to decline. By necropsy, most of the granzyme B expression had returned to baseline levels.

In PBMC samples, neither IFNγ nor IL-10 blockade had an impact on the expansion of NK cells after infection (Fig. S4B). In the BAL, NK cells accumulated more rapidly with IL-10 blockade, i.e., peaking at day 3 post-infection with IL-10 blockade vs. day 7 in the isotype control animals (Fig. S4B). However, there were limited differences in Ki67 expression by NK cells in the BAL between treatment groups, indicating the increase in NK cells at day 3 with IL-10 blockade may be due to enhanced recruitment into the BAL. There were no differences in granzyme B expression in any of the NK subsets between treatment groups (Fig. S4E, F). These data suggest that NK cells in the blood and lungs respond to SARS-CoV-2 infection by upregulating Ki67 and granzyme B production. Moreover, IL-10 may have limited role in regulating the recruitment of NK cells into the lower airways.

Mucosal Associated Invariant T cells (MAIT cells) are innate-like lymphocytes that recognize 5-OP-RU, a small molecule intermediate produced during microbial riboflavin biosynthesis, presented in the context of the MHC-I-like molecule MR1 (*30*). In the context of viral infections, MAIT cells have been shown to be host protective via cytokine-driven, MR1-independent mechanisms (*31-34*). MAIT cells have also been suggested to play a role in SARS-CoV-2 infection (*35, 36*). Therefore, we investigated the MAIT cell responses to SARS-CoV-2 in PBMCs and BAL. MAIT cell frequencies in PBMCs and BAL were relatively stable after SARS-CoV-2 infection (Fig. S5). However, we observed an increase in Ki67 expression by MAITs in PBMCs at day 7 post infection (Fig. S5C). Evidence of Ki67 upregulation without MAIT cell expansion has been reported after MAIT-targeted vaccination of NHP (*37*). At necropsy, we detected MAIT cells in all tissues, with the highest frequency of MAITs found in the BAL. We did not observe any changes in MAIT cell frequency in any compartment with either IL-10 or IFNγ blockade. However, there was an increase in Ki67^+^ MAIT cells in the PBMC and BAL on day 7 in the anti-IL-10 treated animals relative to controls (Fig. S5C). These data suggest that, in rhesus macaques, MAIT cells become activated in response to SARS-CoV-2 but do not significantly expand in frequency. Moreover, IL-10 has a minor role in inhibiting MAIT cell Ki67 expression during SARS-CoV-2 infection.

### B cells and antibody responses

Circulating anti-spike antibodies are correlated with protection against symptomatic SARS-CoV-2 infection provided by vaccines (*38-41*). We next investigated whether IL-10 or IFNγ blockade resulted in changes to B cells and SARS-CoV-2 specific antibody responses. The frequency of total B cells in PBMCs remained unchanged after SARS-CoV-2 infection in all treatment groups (Fig. 3A). In the BAL, there was also no change in the frequency of total B cells after infection in isotype control or anti-IL-10 treatment groups. There was a statistically significant, albeit small, increase in total B cells at day 3 in the rmIFNγR1-Ig treatment group compared to isotype controls. At necropsy, there were no differences in the frequency of total B cells between treatment groups in any of the tissues examined (Fig. 3B). However, germinal center B cells (CD20^+^/BCL-6^+^/Ki67^+^) were decreased in the previously PET-hot pLNs of animals receiving IFNγ blockade (Fig. 3C, D).

Anti-spike and anti-RBD antibodies were detected in the plasma of all animals beginning as early as day 7 post-infection (Fig. 3E). Virus-specific IgM responses peaked on ∼day 14 post-infection and began to plateau or decline by day 28-35. The anti-spike and anti-RBD IgG and IgA responses continued to increase throughout the length of the study and had not yet reached a plateau at the day 35 endpoint. Anti-IL-10 treated animals had higher levels of S-specific IgG and IgA on day 7 relative to the controls. However, by day 14 this difference was no longer apparent, and overall, there were not substantial changes to the S- and RBD-specific antibody responses with either IL-10 or IFNγ blockade.

SARS-CoV-2 neutralizing antibodies were detected in 14 of 15 animals, with the exception of control animal DHMT, which also had the lowest anti-spike and RBD antibody titers at necropsy for all isotypes (Fig. 3F). Plasma anti-RBD IgG and IgA, but not IgM, levels correlated with live-virus neutralization titers at necropsy (Fig. 3G). There were no statistically significant differences in neutralization titers between groups. Altogether, these data show that virus-specific antibody responses to SARS-CoV-2 infection of rhesus macaques are first detectable around day 7 and continue to increase over the first month of infection. While IFNγ may have a role in promoting GC responses in reactive pLNs, it has no major impact on the development of serum antibody responses to SARS-CoV-2 infection. Additionally, while IL-10 may accelerate the induction of serum antibody responses, it did not lead to sustained increases in SARS-CoV-2 specific antibody responses or increases in neutralizing antibody titers.

### Kinetics of SARS-CoV-2 specific T cell responses in the BAL and blood

T cell responses likely have an important role in protection against SARS-CoV-2 infection in humans (*42*). The kinetics of CD4 and CD8 T cells that recognize the major SARS-CoV-2 structural protein antigens of spike (S), nucleocapsid (N), and membrane (M) were measured by *ex vivo* peptide re-stimulation of the PBMCs and BAL cells. Consistent with our previous findings, SARS-CoV-2-specific T cell responses were first detected at day 7 post-infection in both the PBMCs and BAL, S and N-specific T cells were more abundant compared to M-specific T cells, and the frequency of virus-specific T cells in the BAL was ∼5-50 fold higher as compared to the blood (Fig. 4A, B and S6) (*1*). Ki67 expression by SARS-CoV-2-specific T cells peaked at day 7 and rapidly declined to baseline levels by day 28-35 post-infection, suggesting that the turnover of antigen (Ag)-specific T cells declines after the first two weeks of infection (Fig. S7A, B). S-, N-, and M-specific CD4 and CD8 T cell responses were all increased with IL-10 blockade but remained unchanged with IFNγ blockade, as compared to controls (Fig. 4B). The combined S-, N- and M-specific CD4 and CD8 T cell responses were quantified by calculating the area under the curve for each T cell response in each animal (Fig. 4C). The cumulative SARS-CoV-2-specific T cell responses in BAL and PBMCs were significantly increased after IL-10 blockade as compared to controls, suggesting that IL-10 limits early Ag-specific CD4 and CD8 T cell responses to SARS-CoV-2.

**Figure 4.**
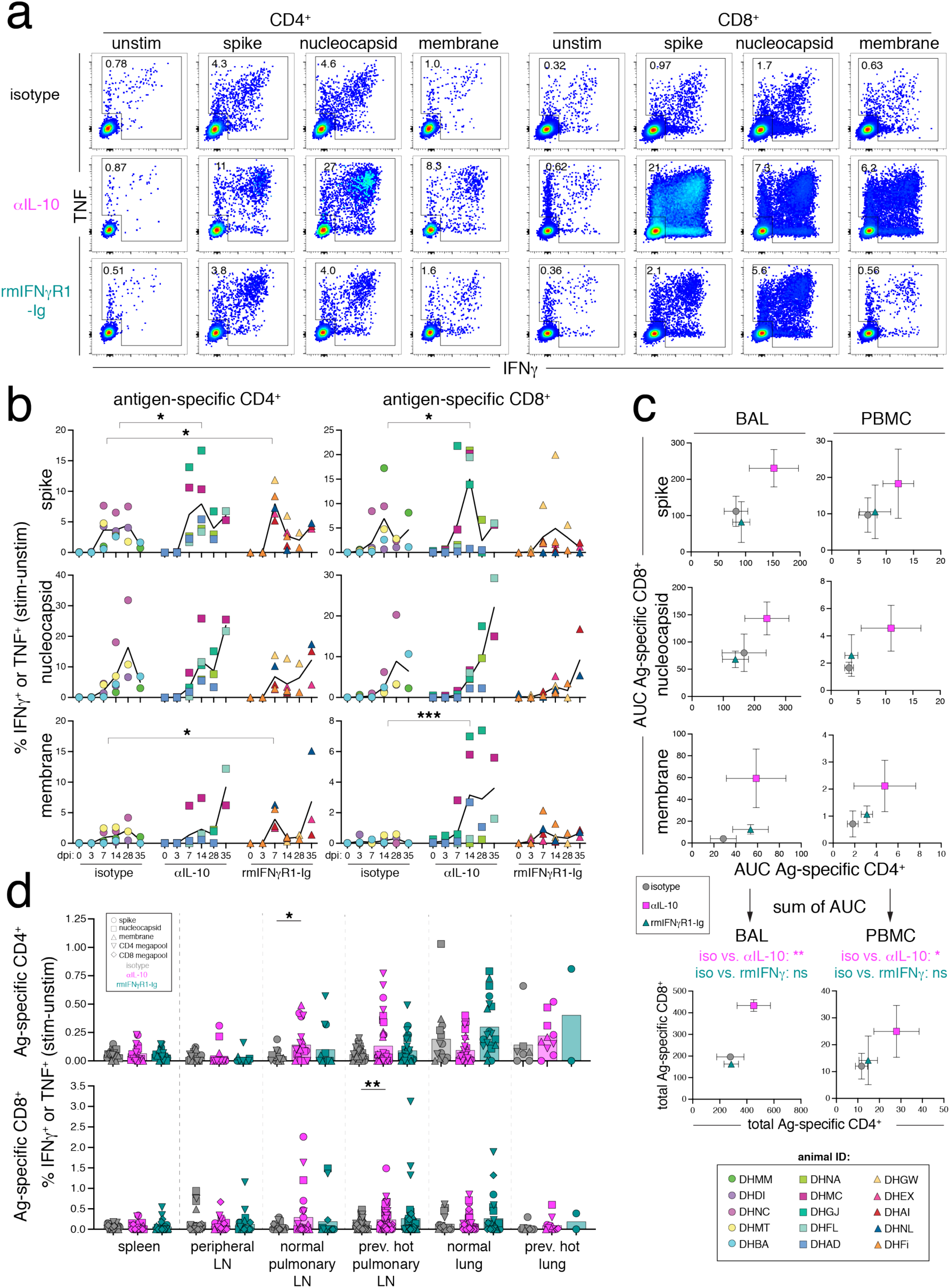
IL-10 blockade increases SARS-CoV-2-specific T cell responses in the blood and BAL fluid. (A) Representative flow cytometry plots of CD4^+^95^+^ and CD8^+^95^+^ T cells from the bronchoalveolar lavage (BAL) at day 14 post-infection responding to *ex vivo* peptide stimulation assay with SARS-CoV-2 15-mer peptide pools for spike (S), nucleocapsid (N), and membrane (M) proteins by production of IFNγ and TNF production. Numbers in plots are the frequency of the gated cytokine+ population. (B) Quantification of frequency of antigen specific CD4^+^95^+^ and CD8^+^95^+^ responses in BAL at baseline (dpi 0), days 3, 7, 14, and necropsy (dpi 28 or 35), calculated by taking the frequency of IFNγ^+^ or TNF^+^ in the stimulated samples and subtracting the frequency in the matched unstimulated samples. Each animal is represented as a point and the mean as a line for each treatment group. Legend is in bottom right corner. Significance calculated by 2-way ANOVA with Dunnett’s multiple comparison test. (C) The mean and SEM of the area under the curve (AUC) for antigen-specific CD4 T cell responses (x-axis) and antigen-specific CD8 T cell responses (y-axis) responses in BAL and PBMC samples calculated from *ex vivo* peptide stimulation with spike (S), nucleocapsid (N), and membrane (M), as represented in *B*. The AUC was determined for dpi 0-28 and interpolated by linear regression for animals necropsied at day 35. The bottom graphs represent the sum of the AUC for the Ag-specific CD4 and CD8 T cell responses, and statistics represent a Dunnett’s multiple comparison test for total AUC responses from treatment groups compared to isotype control. (D) Quantification of frequency of antigen specific CD4^+^95^+^ and CD8^+^95^+^ responses in spleen, peripheral lymph nodes (axillary, cervical, and/or inguinal lymph nodes), normal pulmonary lymph nodes (norm. pulm. LN), previously PET/CT hot pulmonary lymph nodes (prev. hot pulm. LN), normal lung sections (norm. lung), and previously PET/CT hot lung sections (prev. hot lung) at necropsy (dpi 28 or 35), calculated as in *B*. Each animal is represented as a point and the antigen as a shape. Significance calculated by 2-way ANOVA with Dunnett’s multiple comparison test.

T cells responding to S, N, M and SARS-CoV-2 peptide megapools (*43*) were assessed in tissues on day 28-35 post-infection at necropsy. Low frequencies of Ag-specific CD4 and CD8 T cells were detected in the spleen and lymph nodes, with previously hot pLNs having the largest relative responses among the secondary lymphoid organs examined (Fig. 4D). Normal and previously PET-hot lung specimens also had detectable populations of Ag-specific CD4 and CD8 T cells. We observed a slight but statistically significant increase in frequency of Ag-specific T cells in pLNs after anti-IL-10 treatment, with Ag-specific CD4 T cells in normal pLNs and Ag-specific CD8 T cells in previously hot pLNs both being increased compared to isotype control. These data are consistent with the elevated virus-specific T cell responses observed in the BAL after IL-10 blockade and suggest that the effect may have occurred early during priming in lung draining lymph nodes. Furthermore, these data suggest that IFNγ has little role in regulating the expansion of SARS-CoV-2-specific T cells or their migration into the lungs and lower airways.

### Development of tissue-resident memory T cells (T_RM_)

We next examined whether the increase in SARS-CoV-2-specific CD4 and CD8 T cells after IL-10 blockade resulted in other phenotypic and functional changes. The majority of S- and N-specific CD4 and CD8 T cells in the BAL co-expressed the cytotoxic markers granzyme B and CD107a/b (Fig. S7C). In control animals, the proportion of Ag-specific CD8 but not CD4 T cells that expressed granzyme B and CD107a/b increased between days 7 and 14 post-infection. Treatment with rmIFNγR1-Ig resulted in increased granzyme B expression by Ag-specific CD4 T cells at day 7 post-infection and CD8 T cells at day 14 post-infection, as compared to isotype controls (Fig. S7C). Treatment with rmIFNγR1-Ig also resulted in increased CD107a/b expression by both CD4 and CD8 T cells at necropsy compared to controls. Approximately 40% of Ag-specific CD4 T cells co-expressed IL-2, which varied little over time (Fig. S7C). Approximately 10% of Ag-specific CD8 T cells also co-expressed IL-2, which tended to increase at later time points post-infection. There were no differences in IL-2 expression by virus-specific T cells between treatment groups.

Tissue resident memory (T_RM_) T cells have been shown to be important in protection against viral and bacterial infections in the lung and airways (*44-47*). SARS-CoV-2-specific T_RM_ have also been detected in lung autopsy specimens (*48*). We examined expression of T_RM_ markers CD69 and CD103 on S- and N-specific CD4 and CD8 T cells in the BAL. At day 7 post-infection, when virus-specific T cells are first detected, 40-50% of Ag-specific T cells in the BAL express CD69, and <10% of CD8 T cells co-expressed CD69 and CD103 (Fig. 5A-D, S8). At day 14, Ag-specific CD4 and CD8 T cells further upregulate CD69 and 5-20% co-express CD103. At necropsy, most of the SARS-CoV-2-specific T cells in the BAL expressed CD69 and 30-60% co-expressed CD103. N-specific T cells had higher frequencies of T_RM_ markers compared to S-specific T cells (Fig. 5A-D). N-specific CD8 T cells had the highest frequency of CD69 and CD103 co-expression, ∼60% of total, suggesting that nucleocapsid-specific CD8 T cells are highly tissue resident (Fig. 5D, S8). Thus, by ∼one-month post-infection there is a prominent population of SARS-CoV-2-specific CD4 and CD8 T_RM_ in lower airways of the lung.

**Figure 5.**
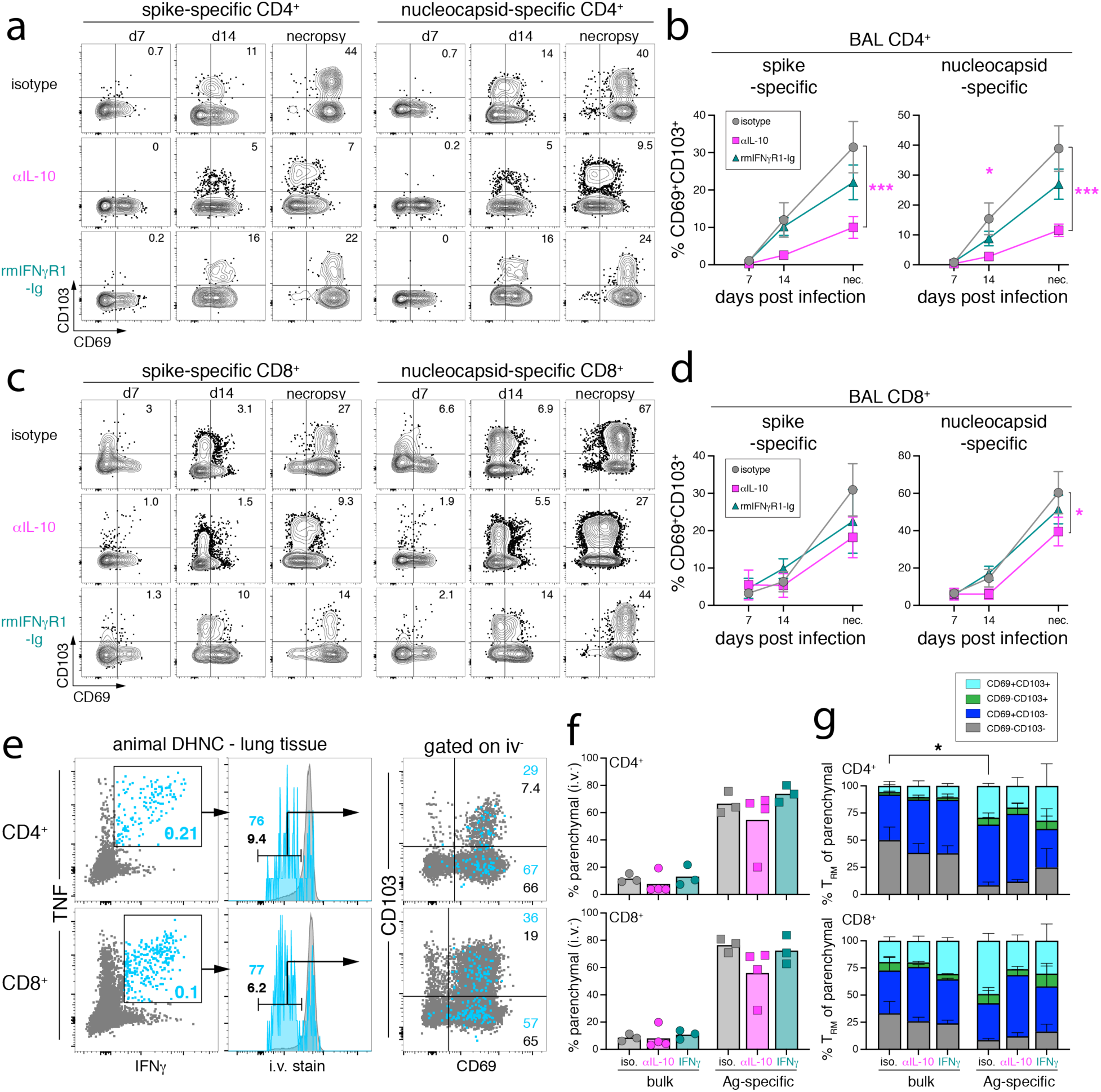
IL-10 blockade impairs the differentiation of SARS-CoV-2-specific T_RM_ cells in the lower airways. (A, C) Representative flow cytometry plots of CD69 and CD103 expression on spike-specific and nucleocapsid-specific CD4^+^ T cells (A) or CD8^+^ T cells (C) from bronchoalveolar lavage (BAL) from day 7, 14, and necropsy (day 28 or 35) post-infection. Numbers in upper right represent the frequency of CD69^+^CD103^+^ gate. (B, D) Quantification of the frequency of CD69^+^CD103^+^ among spike-specific and nucleocapsid-specific CD4^+^ T cells (B) and CD8^+^ T cells (D) from BAL from day 7, 14, and necropsy (day 28 or 35) post-infection. Graph shows mean and SEM. (E) Representative flow cytometry plots of antigen-specific (light blue) and bulk non-specific (grey) CD4^+^95^+^ (top row) and CD8^+^95^+^ (bottom row) T cells from concatenated lung tissue samples from DHNC isotype control animal, at necropsy. Antigen-specific and bulk T cell populations were subsetted for the frequency of intravenous antibody stain (i.v. stain) negative cells, and then among i.v. stain negative cells the frequency of CD69^+^ and CD103^+^. Numbers in gates are the indicated frequencies within the gates. (F) Quantification of the frequency of lung parenchymal localization determined by negative i.v. staining, among bulk and antigen-specific CD4^+^ and CD8^+^ T cells from concatenated lung samples at necropsy, as indicated in *E*. Graphs are plotted as mean and individual animals as points. Significance determined by 2-way ANOVA with Dunnett’s multiple comparison test. (G) Quantification of the frequency of TRM markers: CD69^-^CD103^-^ (grey bars), CD69^+^CD103^-^ (dark blue bars), CD69^-^CD103^+^ (green bars), and CD69^+^CD103^+^ (turquoise bars) among parenchymal antigen-specific and bulk CD4^+^ and CD8^+^ T cells from concatenated lung samples at necropsy, as indicated in *E*. Graphs represent mean and SEM. Significance determined by 2-way ANOVA with Dunnett’s multiple comparison test. For *F* and *G*, only samples with >35 events for antigen-specific cells were used for phenotypic analysis.

Interestingly, IL-10 blockade resulted in a delayed upregulation of T_RM_ markers by SARS-CoV-2-specific CD4 and CD8 T cells, while IFNγ blockade had no effect (Fig. 5A-D). The largest difference in T_RM_ marker expression was observed in the Ag-specific CD4 T cell compartment, with over a 3-fold decrease in frequency of CD69/CD103 double-positive cells among S- and N-specific CD4 T cells in the BAL at necropsy after anti-IL-10 treatment relative to controls (Fig. 5B). We also observed a decrease in CD69/CD103 double-positive T_RM_ among Ag-specific CD8 T cells after IL-10 blockade, but this difference was less pronounced than was observed for CD4 T cells (Fig. 5D). There were no significant differences in T_RM_ frequency with IFNγ blockade.

In the lungs, Ag-specific T cells were preferentially localized to the parenchyma compared to bulk T cells, i.e., very few virus-specific T cells isolated from lung tissue were stained by i.v. delivered antibody (Fig. 5E, F). Of the T cells in the lung parenchyma (i.v.^-^), the majority were T_RM_ and expressed CD69, +/-CD103 (Fig. 5G). More of the Ag-specific T cells in the lung parenchyma were T_RM_ compared to non-specific T cells in the same location (Fig. 5G). This was a significant difference for CD4 T cells, and a similar but non-significant trend was observed for CD8 T cells. Parenchymal localization and T_RM_ marker expression by Ag-specific T cells in the lung did not differ with IL-10 or IFNγ blockade. This contrasted with the observed decrease in Ag-specific T_RM_ in the BAL after IL-10 blockade. Collectively, these data suggest that IL-10 inhibits the expansion of Ag-specific T cells in the lower airways but also promotes their differentiation into T_RM_.

### T cell responses in the nasal mucosa

Immunity in the nasal mucosa is likely critical in protection against SARS-CoV-2 infection (*49*). To detect Ag-specific T cells in the nasal mucosa, we obtained the epithelial lining of the nasal passage (including the nasal turbinates) and restimulated the isolated lymphocytes with the S peptide pool. No S-specific T cells were detected in the nasal mucosa (Fig. 6A-D). This paucity of Ag-specific T cells was not due to an inability to isolate T cells from the nasal mucosa, as intravascular stain-negative, parenchymal T cells with a T_RM_ phenotype (CD69^+^CD103^+/-^) were readily detected (Fig. 6E-H). This is consistent with our previous findings that SARS-CoV-2-specific T cells are not present in the nasal mucosa of rhesus macaques at day 10 post-infection (*1*). Ag-specific T cells were not detectable in the nasal mucosa with either IL-10 or IFNγ blockade, indicating that the lack of virus-specific T cells in the nasal mucosa is not due to IL-10 mediated suppression.

**Figure 6.**
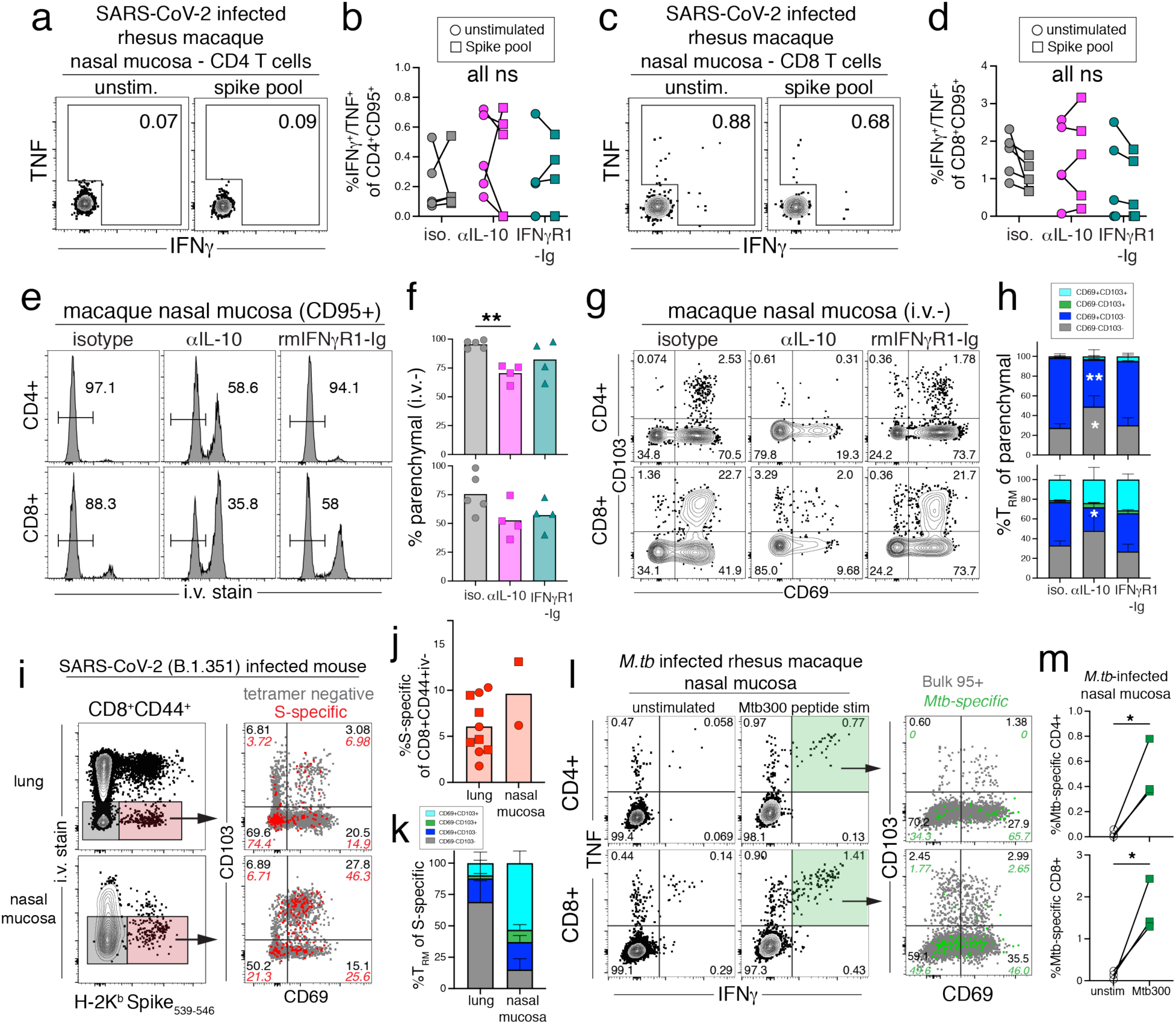
IL-10 blockade does not rescue the lack of SARS-CoV-2-specific T cell responses in the nasal mucosa and erodes pre-existing nasal T_RM_ cells. (A, C) Representative flow cytometry plots of CD4^+^95^+^ (A) or CD8^+^95^+^ (C) T cells responding to spike peptide pool from the nasal mucosa of isotype control rhesus macaque, DHDI, at necropsy (dpi 28) after SARS-CoV-2 infection. Numbers in plots are the frequency of the gated cytokine+ in stimulated or unstimulated samples. (B, D) Quantification of frequency of cytokine+ (IFNγ+ and/or TNF+) CD4^+^95^+^ (B) or CD8^+^95^+^ (D) T cells responding to spike peptide stimulation at necropsy (dpi 28 or 35) form stimulated and unstimulated samples from the nasal mucosa after SARS-CoV-2 infection. Significance calculated with a 2-way ANOVA and Sidak’s multiple comparison test between stimulated and unstimulated samples. (E) Representative flow cytometry plots of bulk CD4^+^95^+^ or CD8^+^95^+^ T cells that stain with intravenous stain (i.v. stain) from the nasal mucosa of rhesus macaques infected with SARS-CoV-2 at necropsy. Numbers in plots indicate the frequency of i.v. stain negative. (F) Quantification of the frequency of i.v. stain negative CD4^+^95^+^ or CD8^+^95^+^ T cells from the nasal mucosa of rhesus macaques infected with SARS-CoV-2 at necropsy (dpi 28 or 35). Significance calculated with 2-way ANOVA and Dunnett’s multiple comparison test. (G) Representative flow cytometry plots of bulk CD4^+^95^+^ or CD8^+^95^+^ T cells expressing CD69 and CD103 from the nasal mucosa of rhesus macaques infected with SARS-CoV-2 at necropsy. Numbers in plots indicate the frequency within the quadrants. (H) Quantification of the frequency of CD69^-^CD103^-^ (grey bars), CD69^+^CD103^-^ (dark blue bars), CD69^-^CD103^+^ (green bars), or CD69^+^CD103^+^ (turquoise bars) of CD4^+^95^+^ or CD8^+^95^+^ T cells from the nasal mucosa of rhesus macaques infected with SARS-CoV-2 at necropsy (dpi 28 or 35). Significance calculated with 2-way ANOVA and Dunnett’s multiple comparison test with isotype control. (I) (*Left*) Representative flow cytometry plots of CD8^+^CD44^+^ T cells staining with i.v. stain and Spike tetramer (H-2K^b^ Spike_539-546_) from the lung and nasal mucosa of mice infected with SARS-CoV-2 (B.1.351) at necropsy (dpi 30). Red shaded gate shows parenchymal (i.v. negative) Spike-specific CD8 T cells and grey shaded gate represents parenchymal (i.v. negative) non-antigen-specific CD8 T cells. (*Right*) Representative flow cytometry plots of CD69 and CD103 expression by parenchymal (i.v. negative) spike-specific (red dots) and non-specific (grey dots) CD8 T cells from the lung and nasal mucosa. Numbers in plots indicate the frequency within the quadrants. (J) Quantification of the frequency of parenchymal (i.v. negative) spike-specific CD8 T cells from the lung and nasal mucosa from mice infected with SARS-CoV-2 (B.1.351) at necropsy (dpi 30). Data shown from two separate experiments, squares represent one experiment and circles represent a second experiment. Nasal mucosa samples were pooled (n=5) prior to staining and each experiment represented as one data point. (K) Quantification of the frequency of CD69^-^ CD103^-^ (grey bars), CD69^+^CD103^-^ (dark blue bars), CD69^-^CD103^+^ (green bars), or CD69^+^CD103^+^ (turquoise bars), of parenchymal spike-specific CD8 T cells from the lungs and nasal mucosa of mice infected with SARS-CoV-2 (B.1.351) at necropsy (dpi 30). (L) (*Left*) Representative flow cytometry plots of CD4^+^95^+^ or CD8^+^95^+^ T cells responding to Mtb300 peptide pool from the nasal mucosa of rhesus macaques infected with *Mycobacterium tuberculosis* at necropsy (14 weeks post-infection). Numbers in plots are the frequency of the quadrants. The green shading represents the frequency of IFNγ^+^TNF^+^ in the Mtb300 stimulated sample. (*Right*) Representative flow cytometry plots of CD69 and CD103 expression by antigen-specific (green dots) and non-specific (grey dots) CD4^+^95^+^ or CD8^+^95^+^ T cells responding to Mtb300 peptide stimulation in the nasal mucosa of rhesus macaques infected with *Mycobacterium tuberculosis*. (M) Quantification of frequency of IFNγ^+^TNF^+^ CD4^+^95^+^ or CD8^+^95^+^ T cells form Mtb300 stimulated and unstimulated samples from the nasal mucosa of rhesus macaques infected with *Mycobacterium tuberculosis*. Significance calculated with individual t-test.

Consistent with the reduction of T_RM_ in the BAL, we observed a decrease in the abundance of T_RM_ in the nasal mucosa with IL-10 blockade (Fig. 6E-H). There was a significant decrease in the frequency of parenchymal CD4 T cells in the nasal mucosa of the anti-IL-10 treated animals compared to controls (Fig. 6E, F). There was also a trend toward a decrease in parenchymal CD8 T cells after IL-10 blockade, but this difference did not reach statistical significance. In the isotype control and rmIFNγR1-Ig treated animals, the majority of CD4 and CD8 T cells in the parenchyma also expressed T_RM_ markers, CD69^+^CD103^+/-^ (Fig. 6G, H). However, the anti-IL-10 treated animals had a significant loss of CD69 expression and a near complete loss of the CD69^+^CD103^+^ subset within the parenchymal T cell subset. Thus, IL-10 not only has a critical role in the formation of SARS-CoV-2-specific T_RM_ cells in the BAL, but it also has a major role in maintaining the bulk population of T_RM_ cells in the nasal mucosa.

We next asked if the absence of SARS-CoV-2-specific T cell responses in the nasal mucosa is unique to rhesus macaques by examining responses in the nasal mucosa of mice. C57BL/6 mice were intranasally infected with SARS-CoV-2 beta strain (B.1.351), and virus-specific CD8 T cell responses in the nasal mucosa and lungs were quantified. At day 30 post-infection, parenchymal K^b^/Spike_539-546_-specific memory CD8 T cells were detected in both the lung and nasal mucosa (Fig. 6I, J). Approximately 50% of the S-specific CD8 T cells in the nasal mucosa were CD69^+^CD103^+^, compared to ∼10% in the lung (Fig. 6K). Therefore, SARS-CoV-2 infection via intranasal inoculation generates Spike-specific T_RM_ in the nasal mucosa of mice, but not rhesus macaques.

Lastly, to ask if rhesus macaques could generate pathogen-specific T cells in the nasal mucosa, we examined Ag-specific T cells in the nasal mucosa of *Mycobacterium tuberculosis* infected animals. Three male rhesus macaques were infected with ∼50 CFU of *Mycobacterium tuberculosis* (Mtb) H37Rv via endobronchial instillation, and Mtb-specific T cells were quantified by intracellular cytokine staining after restimulation with Mtb peptide pools (*50*). Mtb-specific CD4 and CD8 T cells were readily detected in the nasal mucosa of infected rhesus macaques (Fig. 6L, M) and ∼50-65% of the IFNγ^+^TNF^+^ CD4 and CD8 T cells expressed CD69 (Fig. 6L, M). Thus, rhesus macaques generate antigen-specific T_RM_ in the nasal mucosa in response to Mtb infection, and the absence of SARS-CoV-2-specific T cells responses is unlikely due to a technical difficulty in detecting peptide-specific T cells in this tissue.

## DISCUSSION

Here we contrasted the effects of blocking the pro- and anti-inflammatory cytokines IFNγ and IL-10 during SARS-CoV-2 infection. We found that IL-10 limits the intensity and duration of lung lesions observed with ^18^FDG-PET/CT imaging as well as serum markers of inflammation. However, overall disease remained mild, i.e., IL-10 blockade did not lead to life-threatening outcomes or overt symptomology at the dose given in this study. We also found that IFNγ promoted the intensity and duration of lung lesions observed with ^18^FDG-PET/CT imaging, although the effects of IFNγ blockade were less pronounced compared to IL-10 blockade. Despite these changes in pulmonary inflammation, neither cytokine blockade substantially altered the kinetics of viral replication or peak viral RNA loads. This could indicate that SARS-CoV-2 clearance is mediated by cytokines and cell types other than those downstream of IFNγ and IL-10. Alternatively, it may reflect a strong inherent resistance of rhesus macaques to SARS-CoV-2 infection. Nonetheless, these data show that lung inflammation may be modulated during SARS-CoV-2 without the development of severe COVID-19 disease or loss of viral control.

We also examined the role of IFNγ and IL-10 in modulating adaptive immune responses. We found very little impact of IFNγ blockade on any key features of B or T cell responses, apart from a possible reduction in germinal center formation in reactive lymph nodes. It remains possible that myeloid cell function was regulated by IFNγ, but this hypothesis was not evaluated. However, a key finding in this study was the impact of IL-10 on the induction of SARS-CoV-2-specific T cells. We found that IL-10 inhibited the magnitude of virus-specific T cell responses in the circulation, lower airways, and to a lesser extent in pulmonary lymph nodes. The kinetics of Ki67 expression indicated that IL-10 blockade did not prolong the cycling of virus-specific T cells, so the effects of IL-10 in limiting T cell responses may be mediated during the early clonal burst.

Our data also showed a significant role for IL-10 in promoting tissue residency of T cells at mucosal surfaces. We found that IL-10 increases the rate at which SARS-CoV-2-specific T cells differentiated into T_RM_ within the lower airways. These data suggest that IL-10 could play a role in the switch from the proliferative effector stage to the differentiation memory stage of the adaptive immune response. These data are consistent with findings from Thompson et al., which showed that in rhesus macaques IL-10 induces T cells to differentiate into CD103^+^ T_RM_ (*51*). It was hypothesized that IL-10 drives monocytes to increase production of TGFβ, which is known to promote a T_RM_ phenotype (*52, 53*). Rodda et al., demonstrated that SARS-CoV-2-specific CD4 T cells produce IL-10, indicating that T cells themselves could be a source of IL-10 that regulates T_RM_ differentiation (*54*). However, future studies are needed to identify the cellular sources of IL-10 that regulate memory T cell differentiation after SARS-CoV-2 infection.

Despite the robust virus-specific T cell response in the lungs, we failed to detect spike-specific CD4 or CD8 T cell responses in the nasal mucosa of rhesus macaques. This is consistent with our previous findings that spike- and nucleocapsid-specific T cells were not detected in the nasal mucosa on day 10 post infection (*1*). We did, however, detect SARS-CoV-2-specific T_RM_ cells in the nasal mucosa of infected mice, as well as Mtb-specific T cells in the nasal mucosa of *M. tuberculosis* infected macaques. Moreover, it was also recently shown that virus-specific T cells can be detected in the nasal secretions of previously vaccinated humans who acquire breakthrough SARS-CoV-2 infections (*55*). Therefore, it is not clear why primary intranasal SARS-CoV-2 infection fails to induce a primary effector spike-specific T cell response in the nasal mucosa in rhesus macaques. Nonetheless, we found that IL-10 blockade also reduced the number of T_RM_ cells in the nasal mucosa. Taken together with our observations regarding T_RM_ differentiation of virus-specific T cells in the BAL, these data strongly support the hypothesis that IL-10 plays a major role in promoting and/or maintaining the T_RM_ phenotype of T cells at mucosal surfaces.

Macaques do not develop severe COVID-19, and this study was not designed to test therapeutic approaches. However, these data may have implications for the targeting these cytokines in host-directed therapies during SARS-CoV-2 induced pneumonia or as adjuvants during vaccination. It is possible that inflammation mediated by IFNγ can be alleviated by blockade without impairing viral replication. Although our data show that IFΝγ is not required for control of SARS-CoV-2, we cannot rule out the possibility that increasing IFΝγ might lead to enhanced control of viral replication. In fact, there is a clinical trial underway (NCT05054114) assessing the impact of intranasal IFNγ treatment during SARS-CoV-2 infection. Our data also raise the possibility that providing exogenous IL-10 at the time of mucosal vaccination could promote the formation of tissue resident memory T cells. Lastly, we should emphasize that these conclusions are limited to the setting of well-controlled SARS-CoV-2 infection and mild disease, and the role of these cytokines might differ during uncontrolled viral replication or severe COVID-19. Non-human primate models of COVID-19 pneumonia are needed to investigate the cellular and molecular mechanisms of immune-mediated lung damage. Nevertheless, these results provide insight into the regulation of pulmonary inflammation and T cell immunity during effective immune responses to SARS-CoV-2 infection.

## MATERIALS AND METHODS

### Study design

The study was designed to assess the importance of the pro-inflammatory cytokine, IFNγ, and the anti-inflammatory cytokine, IL-10, early during SARS-CoV-2 infection. We set out to measure changes in lung inflammation, viral replication, and cellular immune responses against SARS-CoV-2 infection after cytokine blockade. The study had a predetermined end point of day 28-35 post-infection. The number of animals included in the study was based on previous studies assessing SARS-CoV-2 infection in non-human primates, as well as practical limitations. The Institutional Biosafety Committee approved all work with SARS-CoV-2 in the BSL-3 facility and approved all inactivation methods. All animal experiments were performed under animal safety protocol LPD-25E (rhesus macaque) and LPD-24E (mouse) at the National Institute of Health and approved by Animal Care and Use Committee (ACUC).

### SARS-CoV-2 infection in rhesus macaques

Fifteen, male rhesus macaques, aged 2.5 to 5 years and weighing 3.5-5 kg, were divided into 3 treatment groups (See Figure 1A). The animals were infected/treated in 5 waves of 3 animals per wave. Each wave consisted of one animal per treatment group. Animals were infected with 2×10^6^ TCID_50_ total of SARS-CoV-2/USA-WA-1: 1×10^6^ TCID_50_ in 3mL intratracheally, and 5×10^5^ TCID_50_ in 0.5mL intranasally in each nostril. For all procedures, animals were anesthetized with ketamine and dexmedetomidine. During anesthesia, animals were weighed and monitored for heart rate, respiratory rate, body temperature, and oxygen saturation. Glycopyrrolate and atipamezole were given for recovery from anesthesia. On graphs including data plotted longitudinally, any pre-infection timepoints were represented as day 0.

### SARS-CoV-2 infection in mice

Twelve-week-old female C57BL/6 mice on 45.1 congenic background from Taconic Biosciences were anesthetized with isoflurane and infected intranasally with 6×10^4^ TCID_50_ of SARS-CoV-2 Beta variant (B.1.351) in 30uL of sterile saline. In total, 10 mice were infected in two rounds of five mice each.

### Mycobacterium tuberculosis infection in rhesus macaque

Three male rhesus macaques aged ∼3 years were infected with 30 to 50 CFU of H37Rv strain of *Mycobacterium tuberculosis* (Mtb) diluted in 2mL of sterile saline. Animals were anesthetized with ketamine and dexmedetomidine and Mtb was bronchoscopically instilled in the right lower lobe. Animals were humanly euthanized at a predetermined endpoint of 13-14 weeks after infection.

### Antibody treatment for SARS-CoV-2 infected rhesus macaques

Rhesus macaques were treated with anti-IL-10 antibody [IL-10R1LALA], rhesus interferon gamma receptor-1-Ig Fusion Protein [IFNGR-Ig], or isotype Control antibody rhesus IgG1 (anti-DSP) [DSPR1], i.v. at 10mg/kg at two timepoints, one day prior to infection with SARS-CoV-2/USA/WA-1, and 3 days after infection. All antibodies used in the treatment regimen were engineered and produced by the Nonhuman Primate Reagent Resource (NIH). Anti-IL-10 [IL-10R1LALA] is a rhesus recombinant antibody, with the LALA effector-silenced mutation on IgG1-kappa (RRID AB_2716328, Cat#PR-1517). Rhesus interferon gamma receptor-1-Ig Fusion Protein [IFNGR-Ig], exists as a dimer and is a fusion between rhesus IFNGR1 extracellular domain and CH2-CH3 of rhesus IgG1 with a GGGS linker between IFNGR1 and Fc (RRID AB_2895625, Cat#PR-0021). Control antibody rhesus IgG1 (anti-DSP) [DSPR1] is an anti-desipramine monoclonal antibody recombinant with rhesus IgG1 (RRID AB_2716330, Cat#PR-1117).

### Quantification of in vitro IL-10 and IFNγ signaling with reporter cell lines

HEK-Blue^TM^ IL-10 (InvivoGen Cat#hkb-il10) and IFNγ (InvivoGen Cat#hkb-ifng) reporter cells lines were maintained in selection media per manufacturer instructions and then plated in test media (DMEM + 10% FBS, 1% Pen-Strep) at 50,000 cells per well in 180uL in a flat bottom 96-well plate. 200pg of IL-10 in 10uL was added to each well of HEK-Blue^TM^ IL-10 cells. Increasing 5-fold dilutions of anti-IL-10 (2.5pg-5mg) in 10uL were added to the respective IL-10 containing wells. 40pg of IFNγ in 10uL was added to each well of HEK-Blue^TM^ IFNγ cells. Increasing 2.5-fold dilutions of rmIFNγR1-Ig (2.5pg-5mg) in 10uL were added to the respective IFNγ containing wells. Control wells were plated without blockade reagents or with cytokines. Plates were incubated for 24 hours at 37°C in 5% CO_2_. The following day QUANTI-Blue^TM^ solution for detection was prepared per manufacturers instructions. 180uL of the QUANTI-Blue^TM^ solution was added to a 96-well flat bottom plate. 20uL of the supernatant from the overnight cell culture was added to the QUANTI-Blue^TM^ solution. The mixture was incubated at 37°C in 5% CO_2_ for 2 hours. The amount of cytokine signaling was determined with a color change reaction and quantified with a spectrophotometer at 650nm.

### Blood and BAL collection

Blood and BAL were collected at baseline (5-19 days prior to infection) and on days 3, 7, 14 post-infection, and at necropsy (day 28 or 35 post-infection). Briefly, blood was collected in EDTA tubes and centrifuged at 2,000rpm for 10 minutes at 22°C to isolate plasma. The remaining blood was diluted 1:1 with 1x PBS and added to SepMate™ PBMC Isolation Tubes (StemCell Cat#85450) containing 15mL of 90% Ficoll-Paque density gradient (Cytiva Cat#17144002), and centrifuged at 1,200g for 10 minutes at 22°C. The upper layer was removed and diluted to 50mL with PBS +1% FBS, and then centrifuged at 1,600rpm for 5 minutes at 4°C. The cells were resuspended at 2×10^7^ cell/mL in X-VIVO 15 media + 10% FBS for staining or *ex vivo* stimulation. BAL was collected by instillation of 50mL of pharmaceutical-grade PBS, 10mLs at a time. BAL was then filtered through a 100um filter and centrifuged at 1,600 rpm for 5 minutes at 4°C. The cell pellet was resuspended at 2×10^7^ cell/mL in X-VIVO 15 media + 10% FBS for staining or *ex vivo* stimulation. Complete blood counts (CBC) were measured on an ProCyte DX (IDEXX Laboratories Inc, Westbrook, Maine) using manufacturer instruction and reagents.

### Clinical Measurements

The procedures for clinical observations and measures were described previously (*1*). Blood chemistries were measured at baseline and necropsy. C-Reactive Protein (CRP) was measured at baseline, 2, 3, 7 days post-infection and at necropsy and analyzed in compliance with manufactory instructions for the QuickRead go® instrument with software version of 6.3.6 (AIDIAN Oy, Espoo, Finland) and QuickRead go CRP kit. The system has a range of 5-150 mg/L and values below limit of detection were plotted at 5mg/L. Prothrombin time, activated partial thromboplastin time, Fibrinogen and D-Dimer levels were measured at baseline, 2, 3, 7, 14, 23 days post-infection and necropsy using a Satellite analyzer, kits, and control sets (Diagnostica Stago Inc, Parsippany, NJ). Heart rate, respiratory rate, peripheral capillary oxygen saturation, and temperature were measured with the Avante Waveline Monitor, an implanted temperature transponder (Model IPTT300, Bio Medic Data Systems, Seaford, DE), and a calibrated rectal temperature probe each time the animals were sedated for handling.

### ^18^FDG-PET/CT Acquisition and Data Analysis

Macaques were imaged using a LFER 150 PET/CT scanner (Mediso Inc, Budapest, Hungary) at baseline (6 to 20 days prior to infection), 2, 6, 10 and 22-24 days post-infection time using ^18^FDG (0.5 mCi/Kg) and images of the chest were processed as previously described (*1*). In each chest scan, regions of interest with abnormal density and/or metabolic activity (FDG uptake > 1.5 SUV) were identified as volumes of interest (VOI) or lesions in the day 2 and/or day 6 scan. The VOIs were transferred to the aligned PET/CT images at other time points to assess the change in lesion volume or FDG uptake as previously described. Lesions that appeared to be continuous in the PET/CT but resulted from inflammation in more than one lung lobe (e.g., the right cranial lobe and the right middle lobe) were handled separately at necropsy and were separated in the scan using the branching of the bronchial tree as a guide. This was necessary in 2 instances, once each in DHBA, and DHMC.

In this study, two additional regions were imaged: the upper abdomen including the spleen and transverse colon and the head and neck. Fused PET/CT images of the head and neck were used to estimate (^18^F)-FDG uptake in tonsils (pharyngeal and palatine) and nasal turbinates. Similarly, to regions in the lungs, VOI were drawn on top of the fused PET/CT image for each time point of the study. In this case, only (^18^F)-FDG uptake values were recorded. A single value was recorded for pharyngeal and palatine tonsils by creating a mask that including all of them in one VOI per time point. Both the maxilloturbinates (MT), ethmoturbinates (ET) were included in a single VOI mask per time point. (^18^F)-FDG uptake was also recorded from a region of about 1 mL of the Trapezius muscle to normalize SUV values of tonsils and turbinates across imaging sessions. Fused PET/CT images of the lower thorax and upper abdomen were used to estimate (^18^F)-FDG uptake in the spleen. A single VOI was created on the baseline PET/CT of the subject. Each scan thereafter was then aligned to match that of the baseline PET/CT. The baseline VOI was then copied onto each timepoint to match the volume of the original VOI of the baseline PET/CT. (^18^F)-FDG uptake was recorded from each timepoint within the area of the spleen. (^18^F)-FDG uptake was also recorded for muscle by the same technique as described previously (*56*). The adnominal scan was not available for all subjects, so only those animals where the standard sized VOI could be mapped on all time points are presented in figure 1.

### Necropsy and intravenous stain for SARS-CoV-2 infected rhesus macaques

To assess the parenchyma localization of T cells within the highly vascularized tissue of the lung we utilized the method of intravenous antibody (i.v.) staining prior to euthanasia (*57-59*). Under anesthesia but prior to euthanasia, 10mL of blood was drawn as a negative control. After blood draw, 100ug/kg of anti-CD45-biotin (clone: MB4-6D6, Miltenyi) was infused for 10 minutes prior to human euthanasia. During incubation time, BAL fluid and 60mL of blood was collected. After euthanasia, the lungs and attached airways, nasal turbinates, salivary gland, tonsil, spleen, and lymph nodes (axillary and pulmonary) were prosected. Regions of the lung that were normal or previously had abnormal HU density or FDG uptake by PET/CT analysis were individually excised from the lung, as previously described (*1*). Pulmonary lymph nodes that were normal or previously had FDG uptake above SUV 2.5 by PET/CT analysis were isolated separately. Tissues and lymph nodes were then divided for RNA isolation, fixation and histology, and single cell preparations for flow cytometry. Mtb-infected macaques were humanely euthanized but did not receive i.v. stain prior to necropsy.

### Necropsy and intravenous stain for SARS-CoV-2 infected mice

Prior to euthanasia mice were injected with 2ug i.v. stain anti-mouse CD45 (clone: 30-F11) in 300uL sterile saline. Antibody was left to circulate 3 minutes prior to euthanasia. Lungs and nasal mucosa, including nasal turbinates, were dissected after euthanasia.

### Tissue digestion for rhesus macaque tissue

All tissues were processed into single cell suspensions for *ex vivo* peptide stimulation and flow cytometry, as previously described (*1*). Briefly, tissues were homogenized in gentleMACS C tubes (Miltenyi Cat#130096334). The tonsil, nasal mucosa, and lung were further digested in RPMI + 50U/mL DNase I + 1mg/mL hyaluronidase + 1mg/mL collagenase D (Roche) on a shaker at 220rpm for 45 minutes at 37°C. The digestion reaction was stopped with equal parts PBS + 20% FBS. After homogenization and/or digestion, the salivary gland, tonsil, nasal mucosa, and lung single cell suspensions were filtered through a 100um filter and then centrifuged at 1,600rpm for 5 minutes at 22°C. Cell pellets were resuspended in 40% Percoll® (Sigma Cat# P1644) gradient and centrifuged at 2,000rpm for 20 minutes at 22°C. The spleen and lung were further cleared of red blood cells by resuspending cell pellets in 2mL of ACK Lysing Buffer (Quality Biologicals Cat#118-156-101) for 2 minutes at room temperature, then stopping the reaction with 10-20mL of PBS + 1%FBS. Cells were resuspended at 2×10^7^ cell/mL in RPMI media + 10% FBS for flow cytometry analyses.

### Tissue digestion for mouse tissue

Dissected lungs and nasal mucosa, including nasal turbinates, were placed in 5mL of digest media (RPMI + 50U/mL DNase I + 1mg/mL hyaluronidase + 1mg/mL collagenase D (Roche)) and homogenized in gentleMACS C tubes (Miltenyi Cat#130096334) and then incubated on a shaker at 220rpm for 30 minutes at 37°C. After digestion cells were filtered through a 100um filter and the reaction was stopped with PBS+ 20% FBS. The single cell suspension was then spun at 1,600rpm for 5 minutes. The cell pelleted was resuspended in 5mL of 37% Percoll® (Sigma Cat# P1644) gradient and centrifuged at 2,000rpm for 20 minutes at 22°C without brake. The cell pellet was resuspended in 2mL of ACK Lysing Buffer (Quality Biologicals Cat#118-156-101) for 2 minutes at room temperature, then stopping the reaction with 10mL of PBS + 1%FBS. Cells were resuspended at 2×10^7^ cell/mL in X-VIVO 15 media + 10% FBS for flow cytometry analyses.

### Peptide stimulation assay

Single cell suspensions were plated at 2×10^6^ cells per well in round bottom 96 well plates in 200uL of X-VIVO 15 media, supplemented with 10% FBS, Brefeldin at 1000x (Invitrogen Cat#00-4506-51), Monensin at 1000x (Invitrogen Cat#00-4505-51), CD107a APC at 1:50, CD107b APC at 1:50, and with or without peptide pools at 1ug/mL. Plates were incubated at 37°C + 5% CO_2_ for 6 hours. The spike peptide pool consisted of Peptivator SARS-CoV-2 Prot_S1 (Miltenyi Cat#130-127-048) and Peptivator SARS-CoV-2 Prot_S (Miltenyi Cat#130-127-953). The nucleocapsid peptide pool consisted of Peptivator SARS-CoV-2 Prot_N (Miltenyi Cat# 130-126-699). The membrane peptide pool consisted of Peptivator SARS-CoV-2 Prot_M (Miltenyi Cat# 130-126-703). CD4 megapool consisted of CD4_S_MP and CD4_R_MP, and CD8 megapool consisted of CD8_MP_A and CD8_MP_B, as described (*43*). The frequency of SARS-CoV-2-specific cells was calculated based on the frequency of IFNγ and/or TNF+ cytokine producing CD4 or CD8 T cells in the stimulated wells minus the frequency in the unstimulated wells, as previously described (*1*). The Mtb300 peptide pool consisted of MHC-I and MHC-II peptides from Mtb at 1 and 2ug/mL respectively. The frequency Mtb-specific cells was calculated based on the frequency of IFNγ+/TNF+ cytokine producing CD4 or CD8 T cells. Significance was calculated based on the difference between matched stimulated and unstimulated samples.

### Flow cytometry and antibody staining

Cells for B cell staining panels were resuspended in 50uL Human Fc-Block (BD Cat#564220) diluted to1:500 in PBS + 1%FBS and incubated for 30 minutes at 4°C prior to washing and surface staining. Cells for MAIT cell staining panels were resuspended in 40uL RMPI + Dasatinib at 1000x, Brefeldin at 1000x, and Monsensin at 1000x, and incubated for 10 minutes at 37°C, then 10uL of rhesus macaque MR1 tetramer APC (NIH tetramer core facility) was added and incubated for an additional hour at 37°C. For surface staining, cells were spun down and resuspended in 50uL of antibody cocktail diluted in PBS +1%FBS +10x Brilliant Violet Staining Buffer Plus (catalog#), and incubated at 20 minutes at 4°C. For mouse spike-tetramer staining, H-2K(b) SARS-CoV-2 S_539-546_ (VNFNFNGL) tetramer in APC (NIH tetramer core facility) was added at 1:100 in the surface staining cocktail and incubated for 30 minutes at 4°C.

After surface staining, or between staining steps, cells were washed 3 times in PBS +1%FBS, and resuspended in 100uL of eBioscience Intracellular Fixation & Permeabilization Buffer (Thermo Cat# 88-8824-00) and incubated for 16 hours at 4°C. After fixation, cells were centrifuged at 2,200rpm for 5 minutes at 4°C without brake and washed once with eBioscience Permeabilization Buffer. Cells were resuspended in 50uL intracellular stains diluted in eBioscience Permeabilization Buffer +10x Brilliant Violet Staining Buffer Plus, and stained for 30 minutes at 4°C. After staining cells were washed with eBioscience Permeabilization Buffer 2x and resuspended in PBS + 1% FBS + 0.05% Sodium Azide for analysis on the BD Symphony platform.

### Viral RNA quantification

Nasal and throat swabs were collected at the indicated timepoints (Fig. 1). Swabs were placed in 1mL of Viral Transport Media (1x HBSS, 2% FBS, 100ug/mL Gentamicin, and 0.5ug/mL amphotericin B) and processed as previously described (*1*). Liquid samples, such as swab media, BAL fluid, or plasma, were processed for RNA using using QIAmp Viral RNA mini kit (Qiagen Cat# 52906) and were eluted in 50uL RNase Free water. At necropsy, whole tissues were placed in 1mL RNAlater® media (Sigma Cat# R0901) and stored at 4°C overnight and then stored at −80°C until RNA processing. Tissues were thawed and tissue pieces were transferred to clean 2mL tubes containing 600uL RLT Buffer from the RNeasy RNA Mini kit (Qiagen # 74136) and 2mm beads. Tissues were homogenized with the Precellys Tissue Homogenizer. After homogenization tissue RNA was further processed using the RNeasy Plus Mini kit and RNA eluted in 50uL RNase Free water. Eluted RNA was stored at −80°C long-term.

RT-qPCR reactions for genomic and subgenomic RNA of the N gene of SARS-CoV-2 were performed on all extracted RNA samples, as previously described (*1*). The total reaction volume was 12.5uL for each sample and control, which contained 2.5uL of eluted RNA, 3.25uL Taqpath 1-step RT-qPCR Master Mix (Thermo Cat#A15299), primers at 500nM, probes at 125-200nM, and the remaining volume as RNase free water. N1 genomic RNA was detected with 2019-nCoV RUO Kit, 500 rxn (IDT #10006713), containing CDC 2019-nCoV_N1 Forward Primer (5’-GAC CCC AAA ATC AGC GAA AT-3’), CDC 2019-nCoV_N1 Reverse Primer (5’-TCT GGT TAC TGC CAG TTG AAT CTG-3’), and CDC 2019-nCoV_N1 Probe (5’-[FAM]-ACC CCG CAT TAC GTT TGG TGG ACC-[BHQ1]-3’) at 125nM. N gene subgenomic RNA was detected using Forward Leader sequence primer (5’-CGA TCT CTT GTA GAT CTG TTC TC-3’), sgN Reverse (5′-GGT GAA CCA AGA CGC AGT AT-3’), and sgN Probe (5′-[FAM]-TAA CCA GAA TGG AGA ACG CAG TGG G-[BHQ1]-3′,) at 200nM, all custom made from Eurofins. All samples were tested for RNA integrity using the 2019-nCoV RUO Kit for RNase P, containing CDC RNAse P Forward Primer (5’-AGA TTT GGA CCT GCG AGC G-3’), CDC RNAse P Reverse Primer (5’-GAG CGG CTG TCT CCA CAA GT-3’), and CDC RNAse P Probe (5’-[FAM]-TTC TGA CCT GAA GGC TCT GCG CG-[BHQ]-1-3’). Prepared reactions were read on a QuantStudio 7 Flex Real-Time PCR System, 384-well format (Applied Biosystems Cat# 4485701). Cycling conditions: Initial: 25°C for 2 minutes, 50°C for 15 minutes, and 95°C for 2 minutes, cycling: 95°C for 3 seconds, 60°C for 30 seconds, for 40 cycles. Copies per/mL or copies/gram were calculated based on standard curves generated for each run using an RNA standard of known quantity. The limit of detection was based on the CT limit of detection from the standard curve in each run. For genomic RNA, this was also limited to CT<35, based on manufacturer’s instructions. For subgenomic RNA cutoff CT<37 was used.

### Anti-Spike/RBD Antibody Titers

Anti-SARS-CoV-2 IgG, IgA and IgM antibody titers were quantified with previously frozen plasma samples. Plasma IgG and IgA titers were determined in a dual IgG/IgA assay, with IgG detected by an enhanced chemiluminescent (ECL) based assay and IgA detected by a dissociation-enhanced lanthanide fluorescent immunoassay based on time-resolved fluorescence (DELFIA-TRF, Perkin-Elmer). Specifically, black 96-well plates (PerkinElmer: AAAND-0001) were coated overnight at 4°C with 100uL of Carbonate-Bicarbonate Buffer (Sigma: SRE0034-1L) containing prefusion-stabilized spike (S-2P) at 1ug/mL or RBD at 2ug/mL, as previously described (*60*). The following day, plates were washed with Wash Buffer (1x DPBS +0.1% IGEPAL CA-630 (Sigma catalog 18896-100ml)) using a plate washer and blocked overnight at 4°C with 250uL of Blocking Buffer (1x DPBS +5% dry milk). Plasma samples were thawed, heat-inactivated, and centrifuged at 2,000rpm for 5 minutes to pellet debris, and the supernatant was transferred to a new tube. Plasma was then diluted 1:100 in Dilution Buffer (1x DPBS +5% dry milk +0.2% IGEPAL CA-360). Three-fold serial dilutions were performed across columns in a 96 well plate and 100uL of diluted samples were added to the coated plates. Plates were incubated for 1 hour at room temp on a rotating shaker. After incubation, plates were washed 3 times with a plate washer. 100uL of the secondary antibody mixture consisting of Goat anti-monkey IgG(H+L)-HRP (ThermoFisher: #PA1.8463) at a 1:10000 dilution and Goat anti-monkey IgA-Biotin (Alpha Diagnostic International #70049) at a 1:5,000 dilution in Dilution Buffer was added. Plates were incubated for 1 hour at room temp on a rotating shaker. After incubation, plates were washed 3 times with a plate washer, and 100uL of Streptavidin-Europium (PerkinElmer: 1244-360) diluted 1:2,000 in PBS +0.2% IGEPAL CA-630 was added. Plates were incubated for 1 hour at room temp on a rotating shaker. Plates were washed 3 times with plate washer, and 50uL of HRP substrate (Pierce ECL Substrate, Thermo #32106) was added. After a 10 minute-incubation, HRP-mediated ECL luminescence was read using a Synergy Neo2 HTS plate reader (BioTek) to detect IgG. Then the ECL substrate was removed, and the plates were washed 3 times. After the last wash, 100uL of DELFIA Enhancement Solution (Perkin-Elmer: 4001-0010) was added to generate Europium-based fluorescence. After a 20 minute-incubation, time-resolved fluorescence was read using the Synergy reader to detect IgA. Plasma was analyzed in duplicate plates, and the average reading for each dilution was taken, subtracting the average from blank wells. The cutoff was set at the blank average +3 standard deviations. The titers were determined by interpolating the cutoff on sigmoid standard curve. IgM titers were measured as described for IgA, using secondary goat anti-monkey IgM-biotin (Brookwoodbiomedical,1152) 1:5,000 diluted in Dilution buffer, in a stand-alone DELFIA TRF assay.

### Live virus neutralization assay

Live-virus neutralization assays were performed on Vero-E6 cells stably expressing human TMPRSS2 (Vero-E6T2), as previously described (*1*). One day before the assay, cells were plated in 12-well plates (Falcon Cat#353043) at a density of 0.4 million cells per well in 2mL D10+ medium (DMEM +10% FBS, 1X Glutamax, 1X Anti-Anti (Gibco) and 250ug/mL Hygromycin B (InVivoGen)). The day of the assay, 12-well plates were washed out of selection media with D2 medium (DMEM + 2%FBS, 1X Glutamax). Frozen plasma samples from baseline, says 7, 14, and necropsy (d28-35) post-infection were thawed and inactivated at 56°C for 30 minutes. Samples were then diluted 1:5 and further 2-fold serially diluted. Dilutions were incubated with equivalent volume of 40-50 PFU of SARS-CoV-2 USA-WA1/2020 virus in D2 media for 1 hour at 37°C. After incubating, 100uL of virus/plasma mix + 300uL of D2 media was added to Vero-E6T2 cells, in duplicate. The cells were then incubated for 1 hour at 37°C +5% CO_2_, with occasional gentle agitation. After this incubation, 1.5mL of DMEM +0.6% methylcellulose per well was added. Plates were then incubated for 66-72 hours at 37°C +5% CO_2_. The media was then gently removed and 1mL of Crystal Violet +5% Ethanol and 3% formalin was added, and stained for 20 minutes at room temp. The cell layer was washed with deionized water and then air dried. Plates were scanned and counted for plaques. Percent inhibition was calculated based on plaques at each dilution compared to viral samples incubated without plasma. IC_50_ of percent inhibition was calculated based on non-linear regression.

### Statistical analysis

Data were analyzed using a 2way ANOVA with a Sidak’s multiple comparison test, a Tukey’s multiple comparison test, or a Dunnett’s multiple comparison test, a one-way ANOVA using a Dunnett’s multiple comparison test, or a two-tailed paired t-test. Tests used are indicated in figure legends. For all statistical analysis p<0.05 for the given test is considered significant: * p<0.05, ** p<0.01, *** p<0.001, **** p<0.0001

## ACKNOWLEDGEMENTS

We would sincerely like to thank Danielle E. Dorosky and Nickiana E. Lora for their contributions to the data collection for this study. Blockade reagents used in this study were provided by the Nonhuman Primate Reagent Resource. Funding for this study was provided in part by the Division of Intramural Research/NIAID/NIH. The content of this publication does not necessarily reflect the views or policies of DHHS, nor does the mention of trade names, commercial products, or organizations imply endorsement by the U.S. Government.

