## supplemental data for "IL-10 suppresses T cell expansion while promoting tissue-resident memory cell formation during SARS-CoV-2 infection in rhesus macaques"

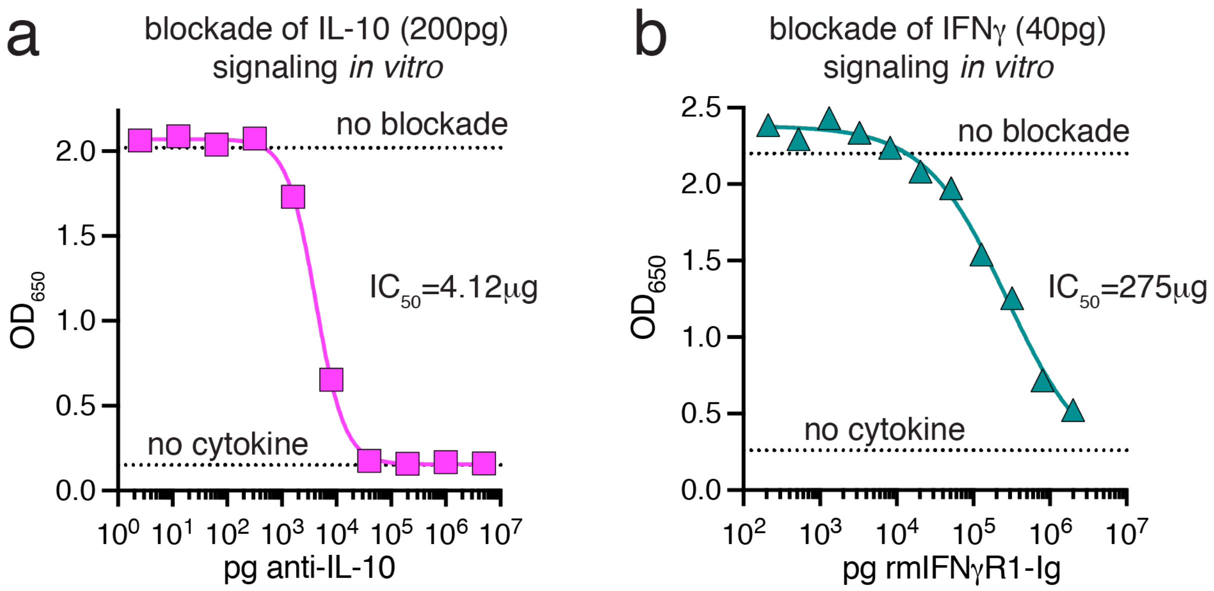


**Supplementary Figure 1. *In vitro* blockade of IL-10 and IFNγ signaling with anti-IL-10 and rmIFNγR1-Ig reagents.** (A) Quantification of the relative amount of *in vitro* IL-10 signaling inhibited by varying amounts of the anti-IL-10 reagent used *in vivo* studies. 200pg of recombinant human IL-10 was added to HEK-Blue™ IL-10 reporter cell line with 5-fold dilutions of anti-IL-10, ranging from 2pg-5mg. The amount of IL-10 signaling was read out with Quanti-Blue solution detection reagent and relative levels enumerated with a spectrophotometer at 650nm. (B) Quantification of relative of the amount of *in vitro* IFNγ signaling inhibited by varying amounts of the rmIFNγR1-Ig reagent used *in vivo* studies. 40pg of recombinant human IFNγ was added to HEK-Blue™ IFNγ reporter cell line with 2.5-fold dilutions of rmIFNγR1-Ig, ranging from 210pg-2mg. The amount of IFNγ signaling was read out with Quanti-Blue solution detection reagent and relative levels enumerated with a spectrophotometer at 650nm. A non-linear sigmoidal fit for *A* and *B* was calculated and the $IC_{50}$ determined for each.

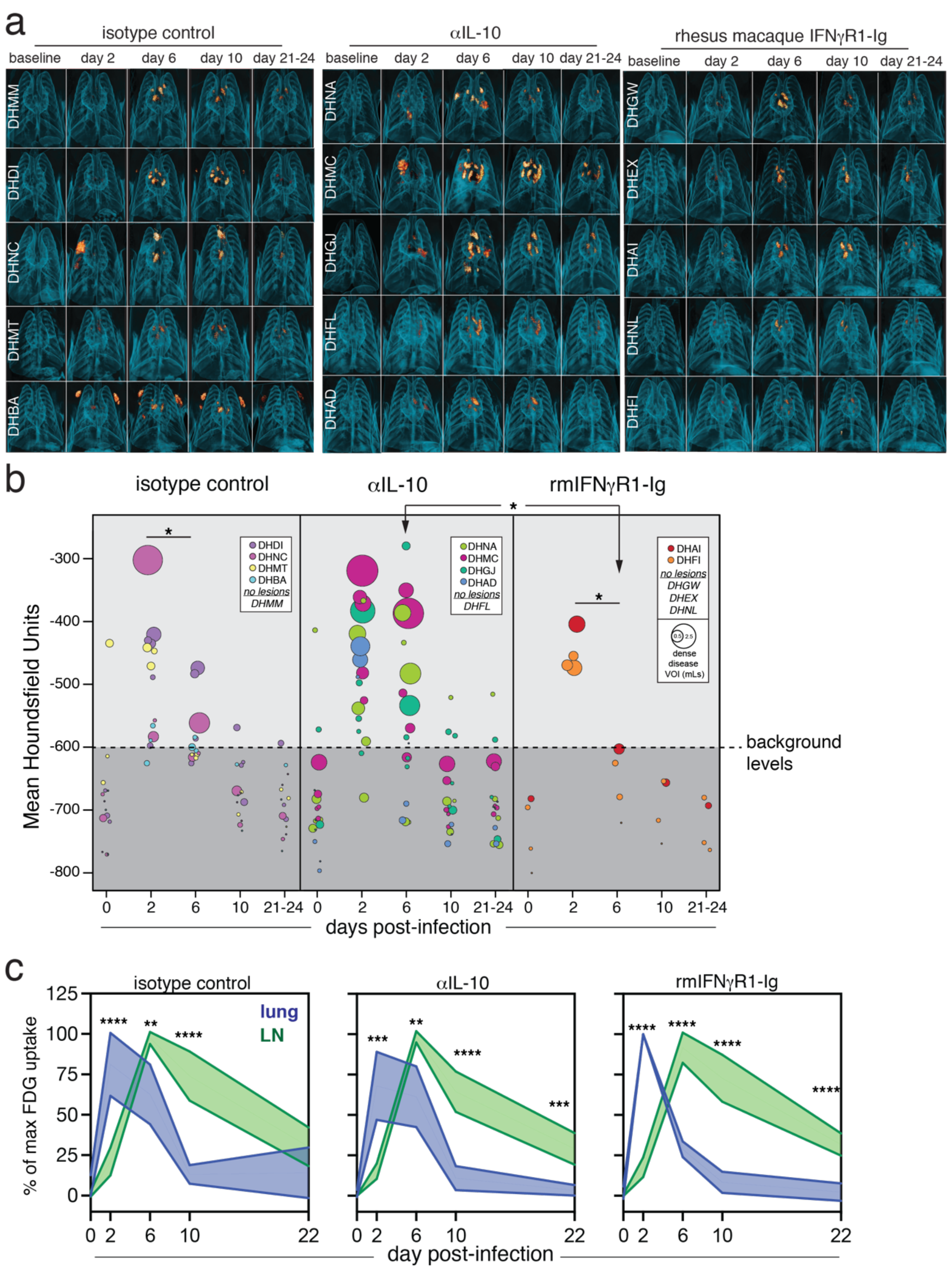


**Supplementary Figure 2. [18]FDG-PET/CT imaging of lungs and lymph nodes of SARS-CoV-2 infected rhesus macaques.** (A) 3D rendering of lung [18]FDG-PET/CT images from baseline, day 2, 6, 10, and 21-24 post infection. Animal IDs are embedded in white. Animals are grouped by treatment. (B) Quantification of mean lung lesion density in Hounsfield Units (HU) (y-axis) and volume of individual lesions (size of dot) over time, based on VOI defined at day 2 or 6 post infection. Significance between groups at each timepoint, and between day 2 and day 6 within groups, was determined by 2-way ANOVA and Tukey's multiple comparison test. (C) Percent of maximum FDG uptake over time in lung and pulmonary lymph nodes plotted a mean with 95% confidence interval. Data normalized for each tissue and each group separately. Significance determined by 2-way ANOVA and Tukey's multiple comparison test between lung and lymph node at each timepoint.

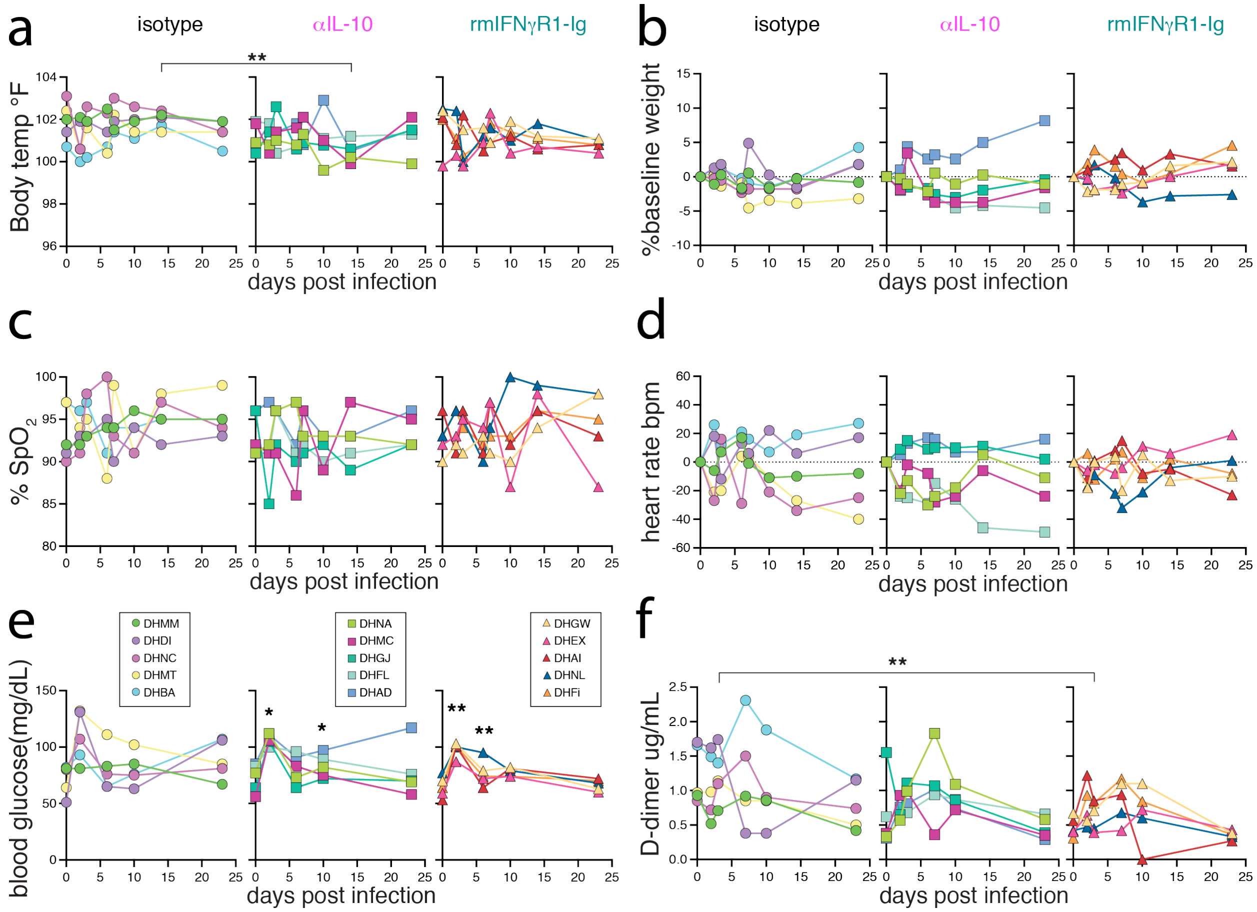


**Supplementary Figure 3. Clinical measurements after SARS-CoV-2 infection.** Laboratory measurements at indicated timepoints after SARS-CoV-2 infection. (A) Body temperature measured rectally in °F. (B) Percent of baseline weight in kg. (C) Oxygen saturation as the fraction of oxygen saturated hemoglobin relative to total hemoglobin in $SpO_2$. (D) Heart rate in beats per minute (bpm). (E) Blood glucose in mg/dL of blood. (F) Plasma D-dimer in ug/mL. Significance calculated by 2-way ANOVA with Dunnett's multiple comparison test.

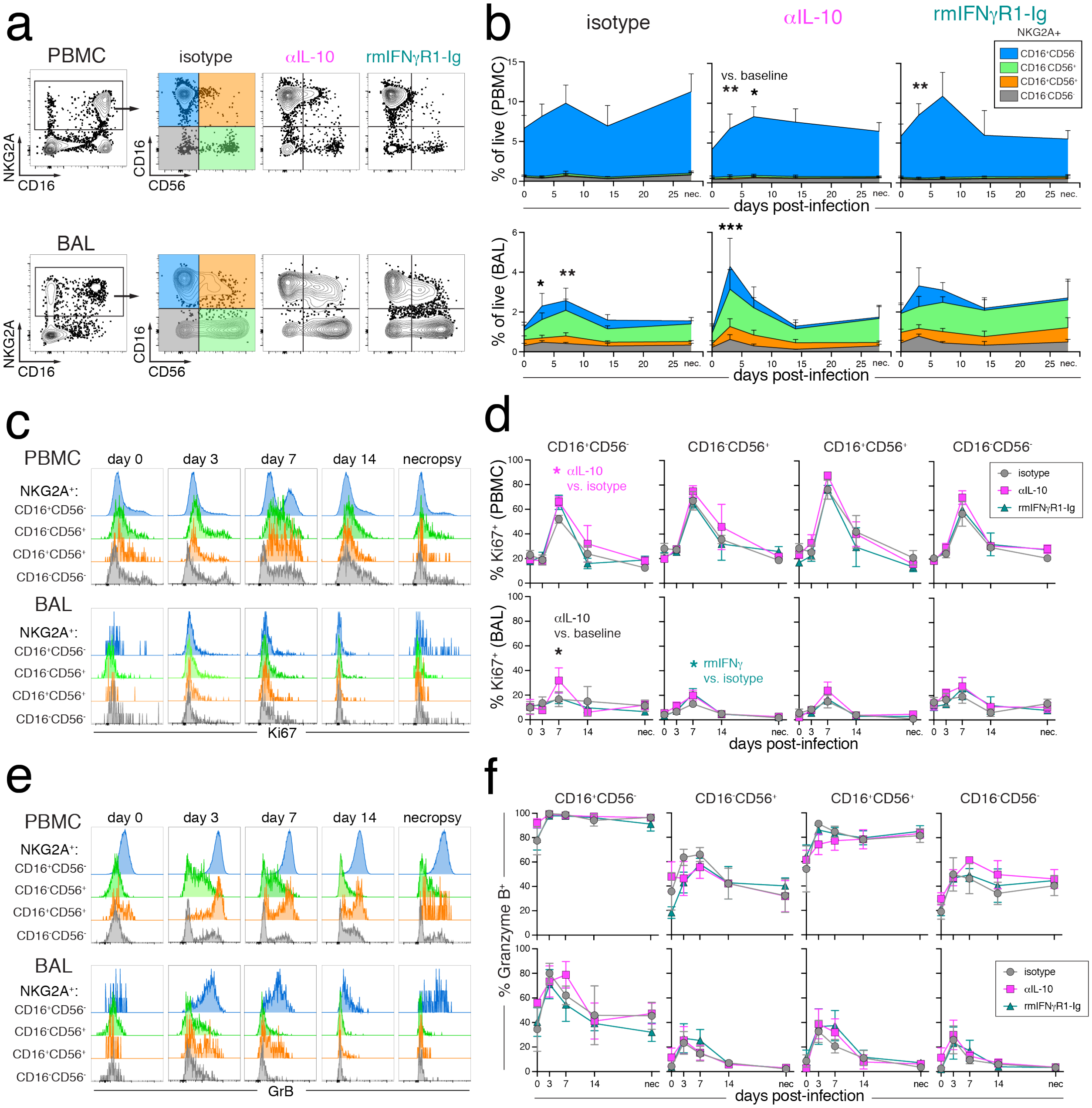


**Supplementary Figure 4. NK cells responses to SARS-CoV-2 infection.** (A) Representative flow cytometry plots of NK cell gating strategy (NKGA2$^+$) and sub-setting (CD56 vs. CD16) in PBMCs and BAL from DHMM (isotype), DHNA (anti-IL-10), and DHGW (rmIFNγR1-Ig) at day 3 after SARS-CoV-2 infection. (B) Quantification of NK cell subset responses as a frequency of total live lymphocytes at baseline, days 3, 7, 14 post infection and necropsy (days 28-35 post-infection). NKG2A$^+$: CD56$^-$/CD16$^+$ (blue), CD56$^+$/CD16$^-$ (green), CD56$^+$/CD16$^+$ (orange), CD56$^-$/CD16$^-$ (grey). Lines are mean and error bars are SEM. Significance calculated with 2-way ANOVA and Dunnett's multiple comparison test of PBMCs at necropsy, isotype control vs. rmIFN$\gamma$R1-Ig; or Tukey's multiple comparison test of BAL at day 3 post-infection as compared to baseline for each group. (C) Representative flow cytometry plots of Ki67 expression by NK cell subsets in PBMCs and BAL from DHMM, DHNA, and DHGW at day 7 post-infection. (D) Quantification of the frequency of Ki67+ of NKG2A$^+$: CD56$^-$/CD16$^+$, CD56$^+$/CD16$^-$, CD56$^+$/CD16$^+$, and CD56$^-$/CD16$^-$ subsets at baseline days 3, 7, 14, and 28/35 post-infection in PBMC (top) and BAL (bottom). Significance calculated with 2-way ANOVA and Dunnett's multiple comparison test. (E) Representative flow cytometry plots of Granzyme B expression by NK cell subsets in PBMCs and BAL from DHMM, DHNA, and DHGW at day 7 post-infection. (F) Quantification of the frequency of Granzyme B+ of NKG2A$^+$: CD56$^-$/CD16$^+$, CD56$^+$/CD16$^-$, CD56$^+$/CD16$^+$, and CD56$^-$/CD16$^-$ subsets at baseline days 3, 7, 14, and 28/35 post-infection in PBMC (top) and BAL (bottom).

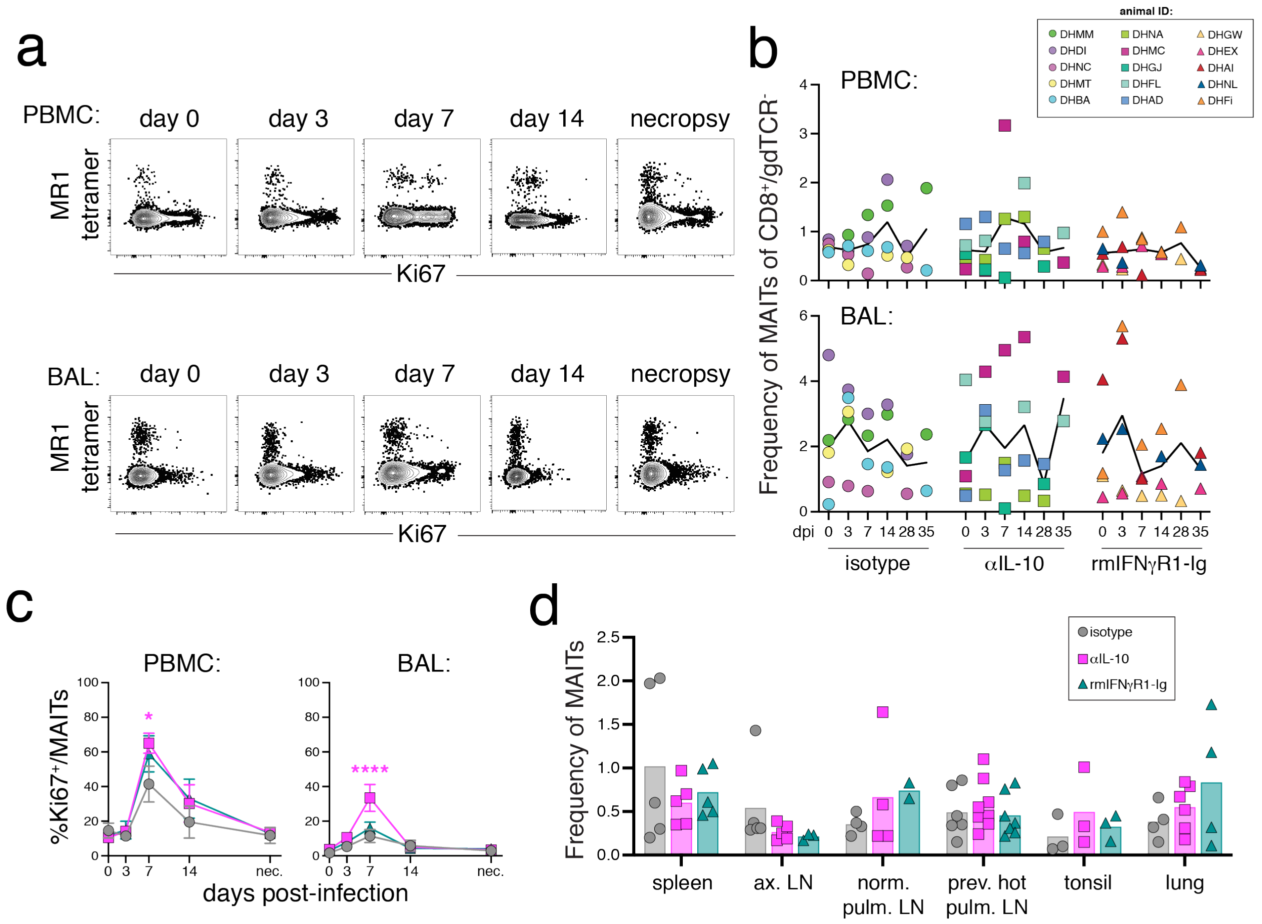


**Supplementary Figure 5. MAIT cell responses to SARS-CoV-2 infection.** (A) Representative flow cytometry plots of MR-1 tetramer staining vs. Ki67 from PBMCs and BAL from DHMM at baseline days 3, 7, 14, and 28/35 post-infection. Plots gated on CD8α$^+$/CD8β$^+$/γδTCR$^-$. (B) Quantification of MAIT cell frequency as percentage of CD8αβ+/γδTCR- in PBMCs and BAL. Each animal is represented as a point and the mean as a line for each treatment group. (C) Quantification of % Ki67+ of MAIT cells in PBMCs and BAL. (D) Quantification of MAIT cell frequency as percentage of CD8αβ+/γδTCR- in the spleen, axillary lymph node (axLN), normal pulmonary lymph node (norm. pulm. LN), previously hot pulmonary lymph node, tonsil, and lung at day 28 or 35 necropsy.

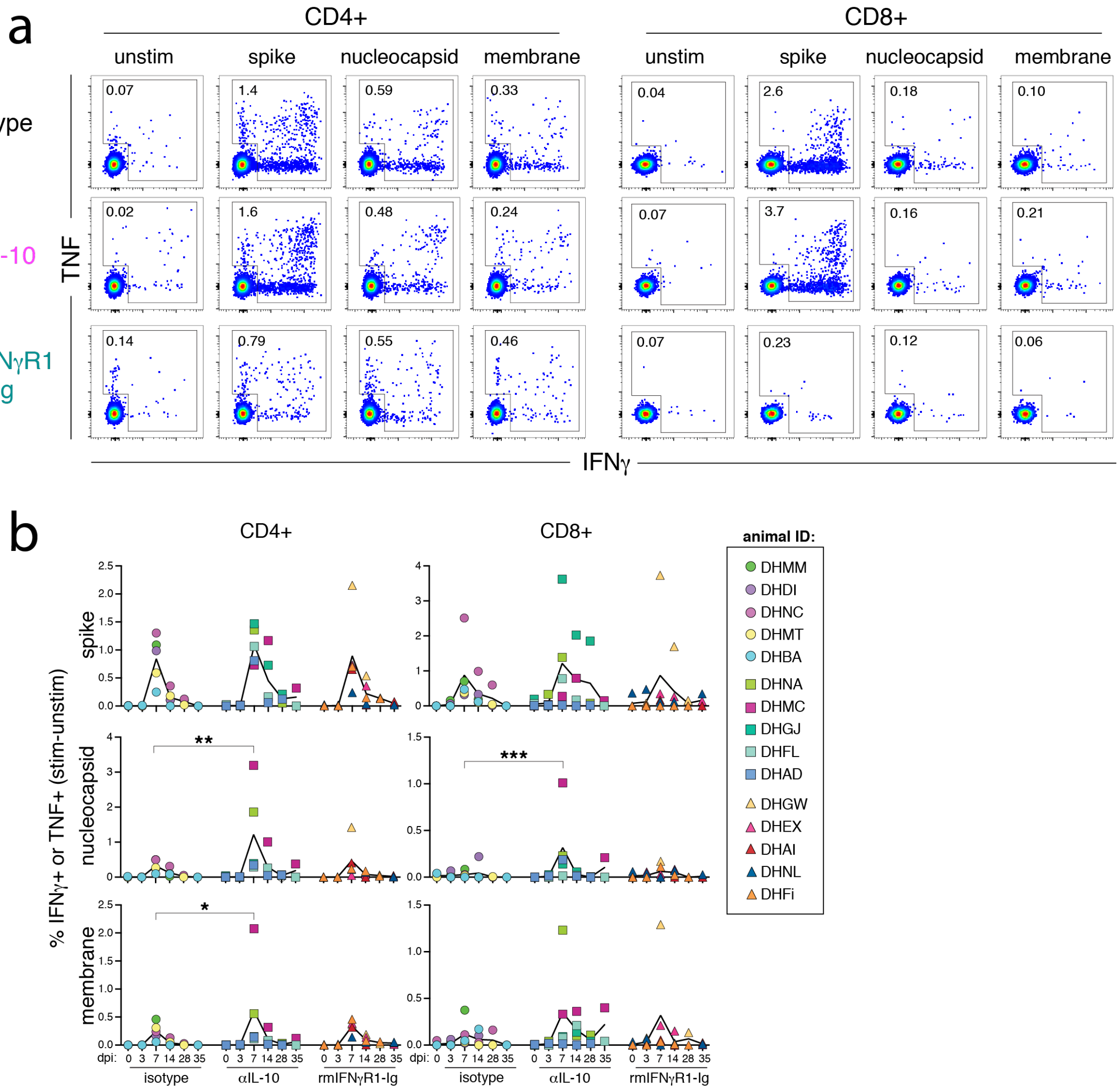


**Supplemental Figure 6. SARS-CoV-2-specific T cell responses in the blood.** (A) Representative flow cytometry plots of CD4$^{+}$95$^{+}$ and CD8$^{+}$95$^{+}$ T cells from PBMCs at day 7 post-infection responding to ex vivo peptide stimulation assay with SARS-CoV-2 15-mer peptide pools for spike, nucleocapsid, and membrane proteins by production of IFNγ and TNF production. Numbers in plots are the frequency of the gated cytokine+ population. (B) Quantification of frequency of antigen specific CD4$^{+}$95$^{+}$ and CD8$^{+}$95$^{+}$ responses in PBMCs at baseline (dpi 0), days 3, 7, 14, and necropsy (dpi 28 or 35), calculated by taking the frequency of IFNγ+ or TNF+ in the stimulated samples and subtracting the frequency in the matched unstimulated samples. Each animal is represented as a point and the mean as a line for each treatment group. Significance calculated by 2-way ANOVA with Dunnett's multiple comparison test at each timepoint between treatment groups and isotype control.

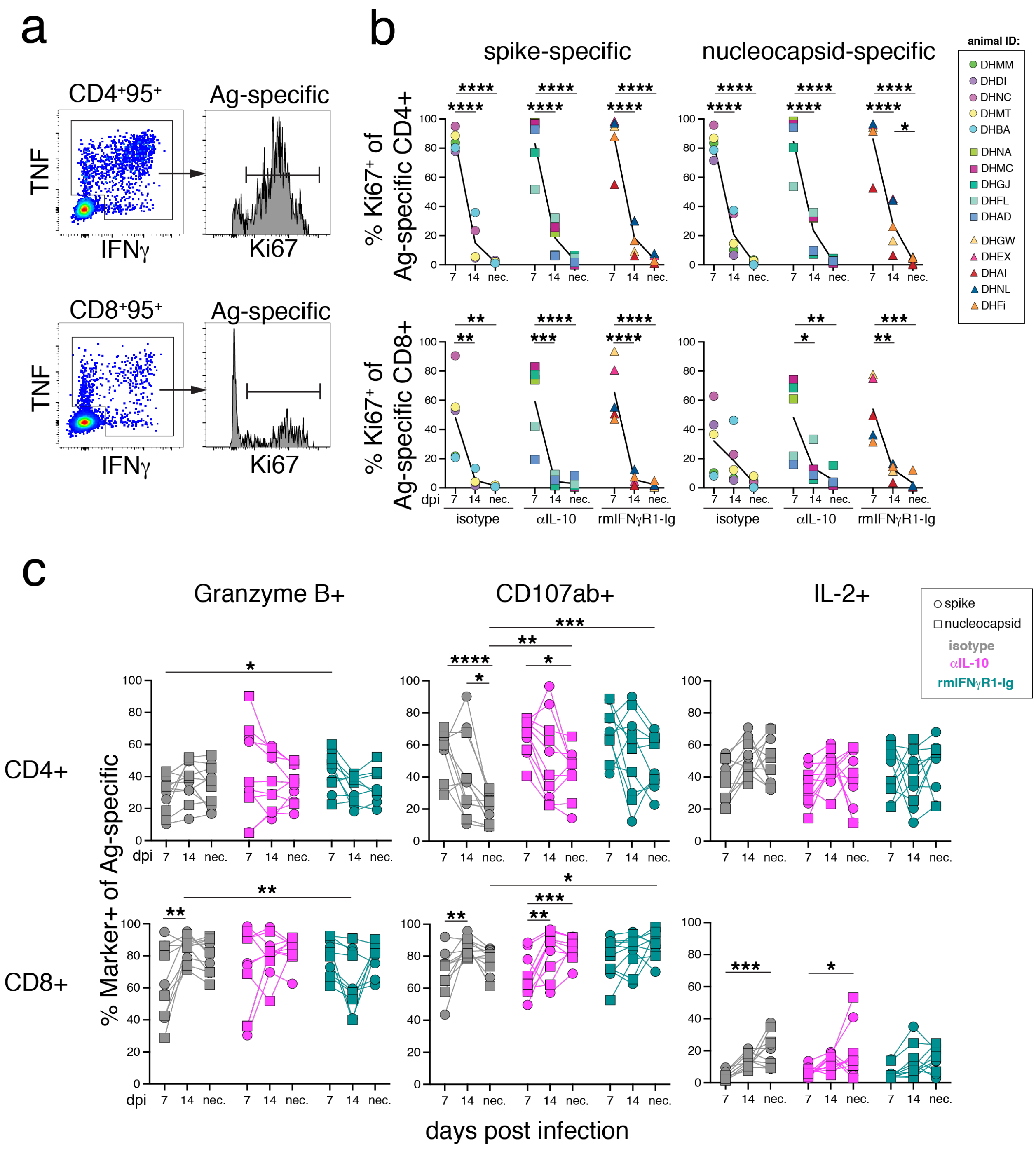


**Supplemental Figure 7. Functionality of SARS-CoV-2-specific T cells.** (A) Representative histograms of Ki67 expression by spike- and nucleocapsid-specific CD4 and CD8 T cells at day 7 post-infection from the indicated treatment groups. CD4 data from group 1 animals. CD8 data from group 4 animals. (B) Frequency of Ki67 expression on spike- or nucleocapsid-specific CD4+95+ or CD8+95+ T cells at day 7, 14, and necropsy (day 28 or 25). (C) Frequency of granzyme B, CD107a/b, and IL-2 expression by antigen-specific CD4+95+ or CD8+95+ T cells at day 7, 14, and necropsy (day 28 or 25). Spike-specific represented as circles and nucleocapsid-specific represented as squares. Significance calculated by 2-way ANOVA with Dunnett's multiple comparison test at each timepoint between treatment groups and isotype control, and within each group comparing changes over time.

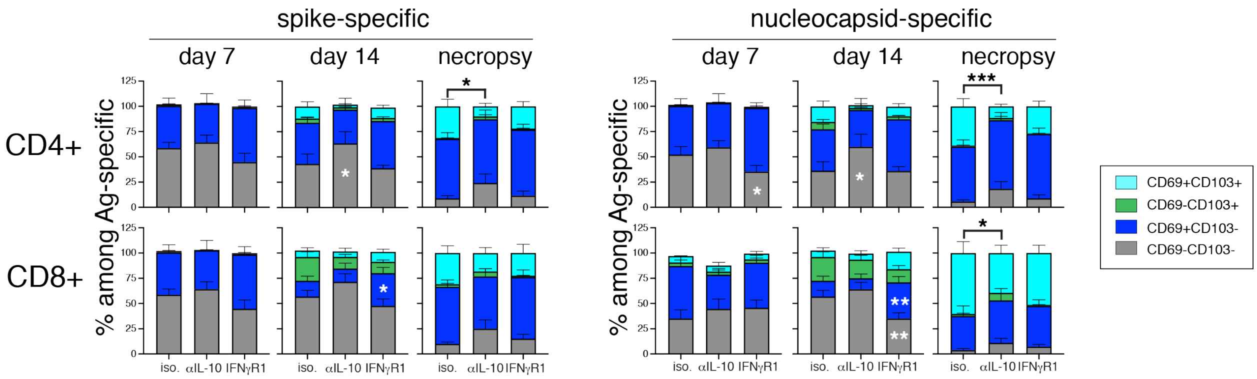


**Supplemental Figure 8. Kinetics of CD69 and CD103 expression on SARS-CoV-2-specific T cells in BAL fluid.** Quantification of the frequency of CD69⁺ or CD103⁺ among spike-specific and nucleocapsid-specific CD4⁺ T cells and CD8⁺ T cells from BAL at day 7, 14, and necropsy (day 28 or 35) post-infection. Graph shows mean and SEM.
